# Robust and scalable inference of cancer progression pathways using Conjunctive Bayesian Networks

**DOI:** 10.1101/2025.07.15.663924

**Authors:** Sayed-Rzgar Hosseini

## Abstract

Cancer is an evolutionary disorder driven by stepwise accumulation of selectively advantageous mutations forming mutational pathways, characterization of which is essential for diagnosis, prognosis and treatment of cancer. Conjunctive Bayesian networks (CBN) are probabilistic graphical models that have enabled the inference of these pathways of cancer progression from genomic data. Previously, we showed that the CBN model can be used to estimate the predictability of cancer evolution as it is able to reflect the underlying cancer fitness landscapes directly from genotypic data. However, the reliability of the inferred pathway probability distributions has not yet been ascertained, which motivates the need for a robust inferential framework. To fill this gap, in this study I have introduced the robust-CBN model (R-CBN). By analyzing synthetic, simulated and real data, I have rigorously compared R-CBN with previous CBN models including CT-CBN, H-CBN, and B-CBN, and the results indicate a superior robustness of the R-CBN model in various settings. Furthermore, I have devised a dynamic programming approximation algorithm, which renders the model amenable to scalability. Thus, R-CBN has the potential to be broadly utilized as a reliable framework to infer cancer-driving evolutionary trajectories, and to distill mechanistic insights from cross-sectional cancer genomic data.

## Introduction

Tumorigenesis is a stepwise process in which accumulation of molecular changes gradually transform normal cells into a malignant neoplasm [1]. This sequence of molecular events can be conceptualized as a pathway along which normal cells progressively gain the ability to evade different layers of growth control mechanisms [2]. Genetic mutations fixed sequentially in a population of cells constitute a prominent example of such pathways [3], and the increasing availability of cross-sectional genomic data, has motivated the development of statistical methods enabling inference of these pathways of cancer progression from mutational data.

Conjunctive Bayesian Networks (CBN) are probabilistic-graphical models [4], which were originally proposed to serve this important purpose and have evolved into different varieties such as CT-CBN [5], H-CBN [6], B-CBN [7] and MC-CBN [8] each addressing different aspects of this challenge. CBN models are particularly well-suited to detect the dependency structure of driver mutations by inferring their partial order of occurrence from cross-sectional mutational data. The assumptions behind the CBN model are well-aligned with the widely held fact that the fitness effect of a mutation depends on the genomic context that is defined by the presence or absence of other mutations [9]. Thus, the dependency structure of driver mutations detected by the CBN model are expected to reflect the underlying fitness landscape on which cancer evolution occurs. In a previous study [10], we showed that this is indeed the case as our CBN-based notion of evolutionary predictability was strongly correlated with the fitness-landscape based one under the Strong-Selection and Weak-Mutation (SSWM) assumption (**Fig. 1**). Despite recent advances in cancer progression modeling, this ability has remained a unique property of the CBN model as this capacity has not been demonstrated for the more recently developed methods, such as Mutual Hazard Networks [11, 12], PMCE [13], HyperTraPS [14, 15] and HyperHMM [16].

**Figure 1.**
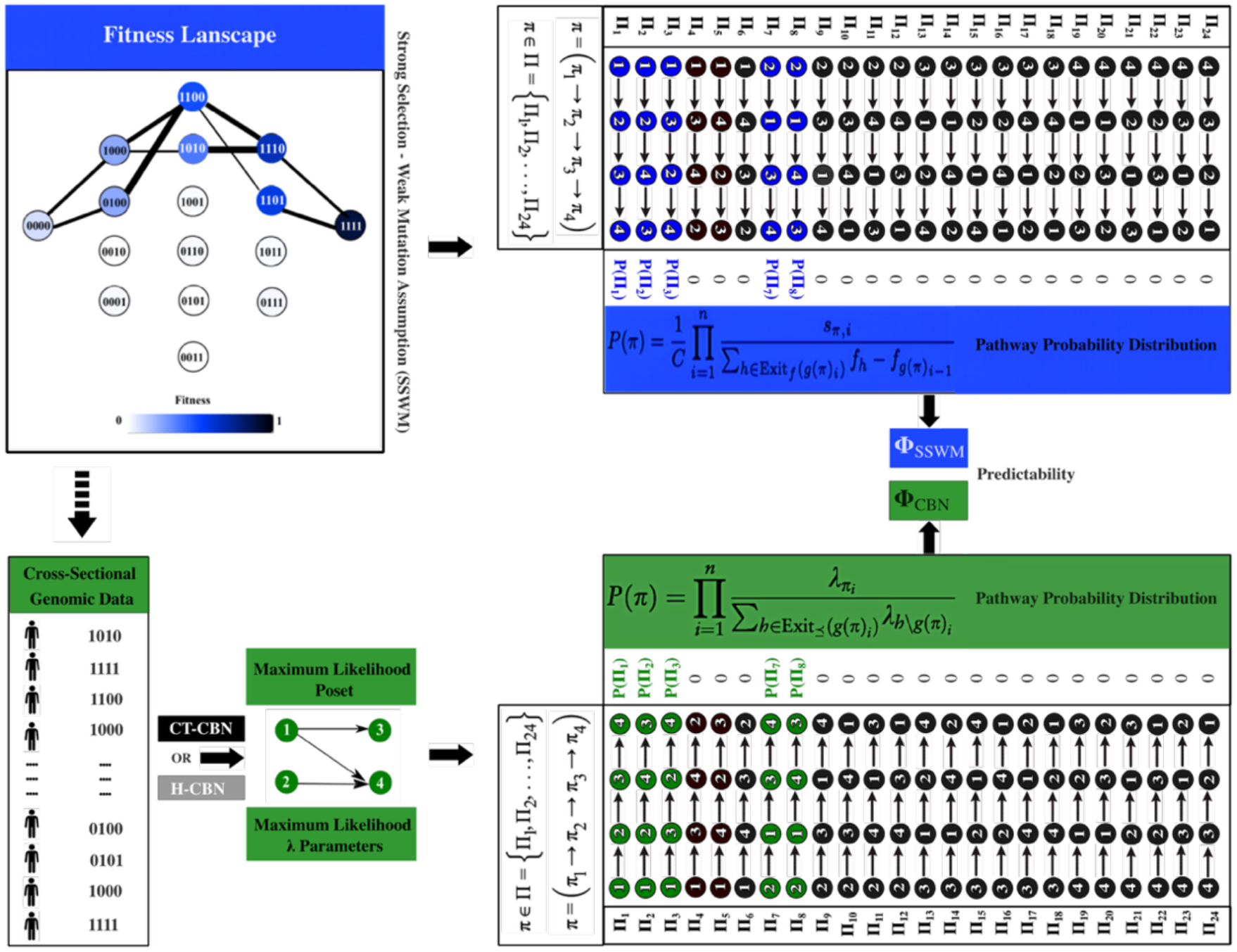
CBN model and the fitness landscapes. Upper part: A fitness landscape is illustrated, which includes 16 binary genotypes that are arranged such that genotypes differing by one single mutation are connected to each other. The fitness assigned to each genotype is color-coded according to the color bar. Under SSWM assumption, along 5 pathways (out of 24) fitness increases monotonically, and the probability of these feasible pathways is computed according to the Eq.4 (See Methods). Lower part: Evolution under the above fitness landscape results in a population of genotypes, which are recorded as cross-sectional genomic data. After training a CT-CBN or an H-CBN model on this data, the maximum-likelihood λ parameters and the poset structure are estimated, which can be used to determine the feasibility of mutational pathways and their corresponding probabilities (See Eq.5 and Eq.6 in Methods). Ideally, the CBN-based probability distribution (green) will be the same as the fitness landscape based one (blue), and the associated predictability quantities (Φ_SSWM_ and Φ_CBN_) computed (using Eq. 21 in Methods) will be equal. Previously, we showed that in various simulation settings, this equivalence almost holds [10].

CBN model enables probabilistic quantification of cancer progression pathways [10, 17], and precise estimation of these probabilities is particularly important. On the one hand, this estimation enables scoring and ranking the vast space of potential pathways, paving the way for mechanistic interpretability. On the other hand, these pathway probabilities can serve as compactly informative quantities that capture the underlying disease context and, therefore, can be used to enable novel pathway-centric disease representation strategies [18]. However, the reliability of these pathway probabilities has been questioned. A previous study has reported large structural variability in repeated evolutionary simulations in fitness landscapes suggesting a many-to-many relationship between the CBN models and fitness landscapes [19]. Further analyses have highlighted that the probabilities estimated by the CBN model may not be reliable [17], which has limited its application to study short-term evolutionary pathways [20]. Thus, there is a critical need for methodological developments to ensure robust and reliable pathway probability estimates, without which broadening the scope of applicability of the CBN model will remain an unattainable goal.

As an initial stride towards this end, in this study I have introduced the Robust-CBN (R-CBN) method. The overarching hypothesis is that relying solely on the maximum-likelihood dependency structure in the CBN model (i.e., the inferred poset or DAG of restrictions) might be the reason behind the non-robust pathway probability estimations. Therefore, the R-CBN algorithm aims to minimize such reliance by exhaustively considering and weighting all potential posets, which are ranked in descending order of their likelihood (**Fig. 2**). However, exhaustive exploration of the space of potential posets quickly becomes prohibitive as the poset size increases, and so this strategy is feasible only for pathways of length four (Quartet R-CBN) or maximally five (Quintet R-CBN). To ensure the scalability of the R-CBN model, I have devised a dynamic programming algorithm (Ensemble R-CBN) that iteratively approximates the probability of a pathway of length *n* from the probabilities of pathways of length *n* − 1. Ultimately, I have rigorously benchmarked the R-CBN method against the previous CBN models including CT-CBN, H-CBN and B-CBN by quantifying pathways of various length using synthetic, simulated and real cancer genomic data.

**Figure 2.**
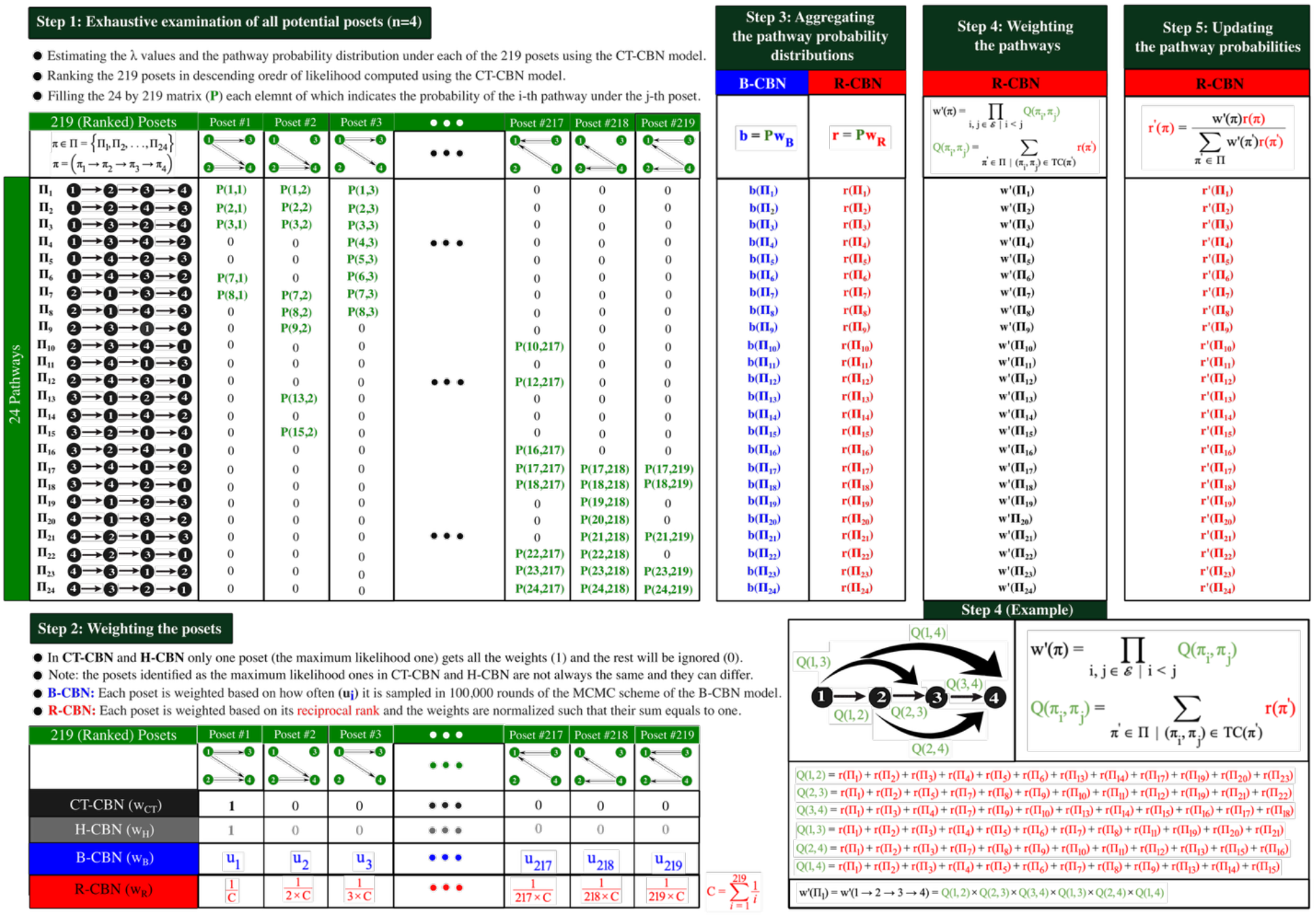
The (Quartet) R-CBN Algorithm. All 219 potential posets of size (*n* = 4) are ranked according to their likelihood, which is calculated using CT-CBN. Then, the probability distribution under each of these 219 posets is determined, which results in a 24 by 219 matrix *P* (step 1). Next, the posets are weighted (*w*_*R*_) based on their reciproal rank (step 2), and then the aggregated pathway probability distribution *r* = *Pw*_*R*_ is quantified (step 3). Similarly, in the B-CBN based algorithm, all the 219 posets of size (*n* = 4) are considered, but their relative ranking is not important as they are weighted (*w*_*B*_) based on how often (*u*) each poset appears in the corresponding 100,000 samples taken using the MCMC sampler. The aggregated pathway probability distribution *b* = *Pw*_*B*_ obtained in step 3 is considered as the final output of the B-CBN based algorithm. In contrast, in R-CBN, *r* is subjected to an additional pathway-level weighting layer (*w*′ defined in step 4) leading to an updated probability distribution (*r*′ in step 5), as the final output. The bottom left panel illustrates how *w*′ in step 4 of the R-CBN algorithm is defined for an example pathway.

## Materials and Methods

### Conjunctive Bayesian Networks (CBN)

CBN models are probabilistic graphical models, which characterize constraints on mutational orders during mutation accumulation processes such as tumorigenesis [4]. A CBN describes the dependency structure of a set ℰ of *n* mutational events as a partial order “*≼*” on ℰ that is represented as a directed acyclic graph (DAG) with vertices ℰ and edges (i, j) for all relations *i* ≺ *j*, such that no *k* ∈ ℰ exists with *i* ≺ *k* ≺ *j* (see an example in **Fig. 1**, bottom).

#### CT-CBN

In continuous-time CBNs (CT-CBNs) [5], the waiting time (*T*_*i*_), for the occurrence of mutation *i* ∈ ℰ is defined recursively as:

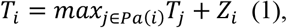

where *Pa*(*i*) denotes the set of parents of *i* in the corresponding DAG, and *Z*_*i*_∼(*λ*_*i*_), *i* = 1, …, *n*, are independent exponentially distributed random variables with rates *λ* = (*λ*_1_, …, *λ*_*n*_). Eq. 1 incorporates the order constraints of ℰ by requiring mutation *i* to occur only after all parent mutations *j* ∈ *Pa*(*i*) have already occurred. Based on the set of observed genotypes *G* as the input, the CT-CBN model ultimately outputs the maximum-likelihood estimates of the λ parameters and the underlying DAG of restrictions using an Expectation-Maximization (EM) algorithm.

#### H-CBN

The hidden CBN (H-CBN) [6] extends the CT-CBN model by additionally allowing observation errors with probability *ε* independently for each mutation, such that the true genotypes become hidden random variables. Based on the set of observed genotypes *G* as the input, the H-CBN model estimates the maximum-likelihood *λ* and *ε* parameters using a nested EM algorithm. To find the maximum-likelihood DAG, H-CBN uses simulated annealing with *T* = 1, which is initiated by the CT-CBN-based maximum-likelihood DAG as the starting poset.

### Mutational pathways

Considering a set ℰ of *n* mutational events, any genotype *g* as a subset of ℰ can be defined as a binary string representing the occurrences of all mutations in *g* ⊂ ℰ. For example, considering *n* = 4 mutations, a genotype with *g* ={2,4} mutations, is shown as 0101. In this framework, a mutational pathway *π* of length *n* in *U* = {0,1}^*n*^ is a permutation *π* = (*π*_1_ → *π*_2_ → ⋯ → *π*_*n*_) ∈ Π, indicating the temporal order of the mutations. Note that Π = *S*_*n*_ is the set of all *n*! permutations of ℰ, which includes all potential pathways of length *n*. Equivalently, the mutational pathway *π* can be denoted as an ordered list of *n* + 1 genotypes *g*(*π*) = (*g*_0_, *g*_1_, …, *g*_*n*_), where 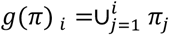, which includes the genotypes successively accumulating the mutations in *π*. For example, in **Fig. 1** the mutational pathway Π_7_ = (2 → 1 → 4 → 3) ∈ Π can be represented by the ordered list of genotypes (0000, 0100, 1100, 1101, 1111).

### Inference of pathway probability distributions

#### fitness landscape-based model

As we described previously [10], the probability of a mutational pathway could be quantified under the fitness-landscape-centered evolutionary models using Strong Selection Weak Mutation (SSWM) assumption [21, 22]. Under SSWM regime, mutations are fixed sequentially in a population, resulting in multi-step evolutionary pathways, whose probability is quantified as the product of fixation probability of mutations, which is proportional to the selective coefficients of the mutations [23]. Under a given fitness landscape *f*, to every potential genotype *g* a fitness value *f*_*g*_ is assigned, and for a given mutational pathway *π*, the selective coefficient *s* of the mutation *π*_*i*_ is defined as the fitness difference that it cases along the mutational pathway:

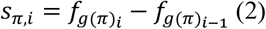

A mutational pathway under the SSWM assumption is feasible, only if the fitness of genotypes is monotonically increasing along the pathway. Hence, the set of all feasible pathways of length *n* is defined as:

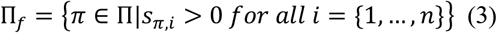

and the probability of a given mutational pathway *π* of length *n* is determined as:

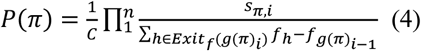

if *π* ∈ Π_*f*_ and zero otherwise [24], where C is a normalizing constant used to ensure that *P*(Π) is a probability distribution (i.e., 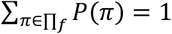).

#### CBN-based model

As we have described previously [10], after training a CBN model (ℰ,*≼*) with waiting time parameters λ, the probability of mutational pathways can be quantified, first by identifying the feasible mutational pathways, which are the linear extensions of the maximum-likelihood poset:

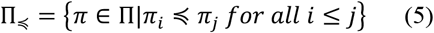

Then, the probability of a given pathway π of length *n* is computed as the product of competing exponentials by recording all possible one-step extensions of *g*(*π*)_*i*_ in its exit set at each step *i*:

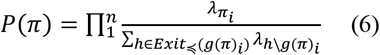

if *π* ∈ Π_*≼*_ and zero otherwise.

### Robust inference of pathway probability distributions (Quartet R-CBN and B-CBN)

The main idea behind the R-CBN algorithm is to reduce the reliance on the maximum-likelihood poset when computing the pathway probabilities by considering and weighting all the potential posets. As an alternative strategy, I have also considered B-CBN approach, which enables weighting the posets using an MCMC-sampling technique [7]. The steps of both algorithms are depicted in **Fig. 2**, and are delineated below:

#### Step 1: Exhaustive examination of all potential posets

R-CBN examines all unique posets represented by the set of transitively closed DAGs of restrictions between a given number (*n*) of mutations. Note that the number of all unique posets quickly grows by increasing *n*; at *n* = 4 (quartets) it is 219, and at *n* = 5 (quintets) it is 4231, but at *n* ≥ 6 examining all unique posets becomes computationally infeasible. Although the following explanation of the algorithm is specific to quartets (*n* = 4), the same procedure can be implemented for *n* = 5 albeit it is computationally more demanding. Under each poset, the CT-CBN is executed and the posets are ranked in descending order of their likelihood calculated by the CT-CBN model. Then, based on the maximum-likelihood estimation of the *λ* values under each poset, the probability *P*(*π*) of each of the *n*! potential pathways *π* ∈ Π, are calculated using Eq. 6. After this step, for *n* = 4, a 24 by 219 matrix *P* will be filled, each element *P*(*i, j*) of which represents the probability of the *i*^*th*^ pathway under the *j*^*th*^ poset. It is important to note that in the B-CBN-based method the matrix *P* is also used, but B-CBN does not require the posets to be ranked.

#### Step 2: Weighting the posets

In the CT-CBN and H-CBN methods, the focus is entirely on the maximum-likelihood poset and so the other 218 potential posets are neglected when calculating the pathway probabilities. However, in R-CBN and B-CBN all posets are considered, but they are weighted differently. In R-CBN posets are weighted (*w*_*R*_) based on their reciprocal rank, which is a common unsupervised strategy for fusing multiple individual ranking methods [25]. In other words, the weight assigned to the *i*^*th*^ ranked poset is 1/(*i* × *C*) where 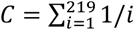. In the B-CBN based approach, however, the relative ranking of the posets is not considered, but rather posets are weighted (*w*_*B*_) according to their relative frequency (*u*_*i*_) in the samples taken by the MCMC sampling strategy.

#### Step 3: Aggregating the pathway probability distributions

Using the weights defined in step 2, the 219 probability distributions in the P matrix (step 1) are aggregated as 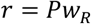 in R-CBN and 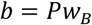 in B-CBN. Both *r* and *b* are vectors of length 24 each element of which corresponds to one of the 24 pathways of length 4 in this example. It is important to note that the vector *b* is the final pathway probability distribution outputted by the B-CBN approach, whereas the vector *r* in R-CBN requires further modifications in steps 4 and 5.

#### Step 4: Weighting the pathways

The above weighting scheme moderates the pathway probabilities as the aggregated probability distribution can largely level out, and consequently the predictability can be substantially compromised. Nevertheless, the information hidden in the likelihood-based ranking of the posets is rich enough to enable R-CBN to offer an intrinsic solution to overcome this limitation. R-CBN can self-update the pathway probabilities by leveraging the edge probabilities, which are the common elements in the structure of both posets and pathways. The main idea is that if there exists a constraint that implies mutation *i* to occur before mutation *j*, the posets having the edge (*i, j*) as a part of the transitive closure of their corresponding DAG, will be ranked higher in terms of likelihood than the posets having the opposite edge (*j, i*). Consequently, the sum of the probability of the pathways in which *i* occurs before *j* (i.e., the marginal probability of the (*i, j*) edge denoted as *Q*(*i, j*)), will be higher than that of the opposite edge (*j, i*) denoted as *Q*(*j, i*). R-CBN utilizes these edge probabilities *Q* to derive the pathway weights (*w*′) as follows:

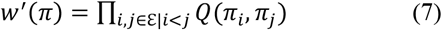

where *Q*(*π*_i_, *π*_j_) is the marginal probability of the edge (*i, j*) in the pathway *π*, which is defined as follows:

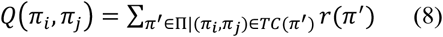

where *TC*(*π*′) is the transitive closure of the pathway *π*′. As the example in Fig. 2 illustrates, in the case of *n* = 4, the multiplication in Eq. 1 involves the probabilities of the 6 edges in the transitive closure of the given pathway. Furthermore, the summation in Eq. 2 involves the probabilities of half of the pathways (12 out of 24), which are compatible with the given edge.

In summary, the pathway-level weighting (*w*′ in step 4) is intended to complement the poset-level weighting (*w* in step 2) to get most out of the information hidden in the relative ranking of the posets (in step 1) in order to update the pathway probabilities (r in step 3), such that the final probability distribution (*r*′ in step 5) more closely reflects the existing constraints in the temporal ordering of mutations.

#### Step 5: Updating the pathway probability distributions

Using the *w*′ weight vector, the updated pathway probability distribution (*r*′), which is the final output of the algorithm, is derived from *r* as follows:

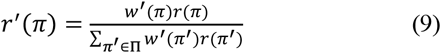

### Scalable inference of pathway probability distributions (Ensemble R-CBN)

Exact inference of the probability of pathways of length (*n* ≥ 6) using the R-CBN approach is not feasible as exhaustive exploration of all potential posets quickly becomes prohibitive at larger poset sizes (*n* ≥ 6), which requires an approximate inferential scheme. For this purpose, I have designed a dynamic programming algorithm (Ensemble R-CBN) that enables inference of the probability of pathways of any given length (*n* > 4) through a bottom-up approach starting from pathways of length *n* = 4 inferred by the Quartet R-CBN. The Ensemble R-CBN algorithm is founded on the following theorem, which enables approximation of the probability of a given pathway of length *n* from the probabilities of pathways of length *n* − 1.

#### Theorem 1

The probability of a given pathway *π* of length *n*, which is a permutation of a set (ℰ) of *n* mutational events, can be approximated as follows:

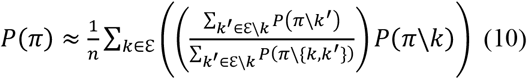

***Proof*:** A pathway *π* of length *n* can be obtained by inserting a new mutation *k* into a specific position along its sub-pathway *π*′ = *π*\*k*, which is of length *n* − 1. It is important to note that there are *n* − 1 other potential pathways of length *n* that can be obtained by inserting the new mutation *k* into alternative positions along the pathway *π*′. Let’s define 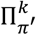 as the set of *n* pathways of length *n* that can be obtained by inserting the new mutation *k* into one of the *n* potential positions along the pathway *π*′: 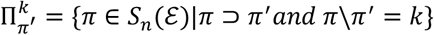 (11), where ⊃ denotes the “ordered superset” symbol. By summing up the probabilities of the pathways belonging to 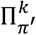 we basically marginalize over the “position of the mutation *k*”. Therefore, we have 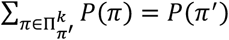 (12). It can be equivalently stated that *P*(*π*′) is partitioned among these *n* pathways of length *n*, so the probability of each 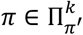 is a fraction (*α*_*π*_′_,*π*_) of *P*(*π*′):

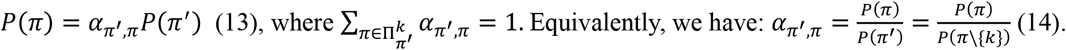

The fraction (*α*_*π*_′_,*π*_) can be intuited as the relative propensity of the mutation *k* to the specific position in *π*′ = *π*\*k*, where insertion of mutation *k* results in pathway *π*, as compared to the alternative positions in *π*′, where insertion of mutation *k* results in alternative pathways in 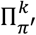. The main assumption is that the insertion propensity of the mutation *k* to a given position in *π*′ on average will remain approximately the same for all sub-pathways of *π*′. For example, if we iterate one step backward by removing a given mutation *k*′ ∈ ε\*k* from both *π*′ and *π*, we obtain their corresponding sub-pathways *π*′\*k*′ and *π*\*k*′, which are of length *n* − 2 and *n* − 1 respectively. The main assumption is that the insertion propensity (*α*_*π*_′_\*k*_′_,*π*\*k*_′) of the mutation *k* to the specific position along *π*′\*k*′ that results in *π*\*k*′, on average will remain equal to the insertion propensity (*α*_*π*_′_,*π*_) of mutation *k* to the position along *π*′ that results in *π*. Thus, we assume the weighted average of *α*_*π*_′_\*k*_′_,*π*\*k*_′ over all *k*′ ∈ ε\*k* remains approximately the same as *α*_*π*_′_,*π*_:

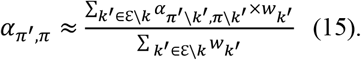

According to Eq. 14, we will have:

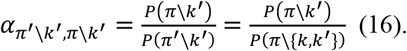

Replacing *α*_*π*_′_\*k*_′_,*π*\*k*_′ in Eq. 15 by Eq. 16 leads to:

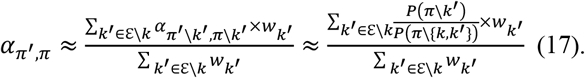

By weighting each *k*′ based on the probability of their corresponding *π*′\*k*′ pathway (i.e., *w*_*k*_′ = *P*(*π*′\*k*′) = *P*(*π*\{*k, k*′}), we can simplify Eq. 17 as follows:

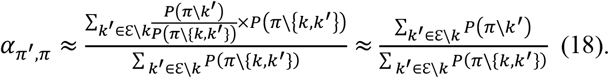

Replacing *α*_*π*_′_,*π*_ in Eq. 13 by Eq. 18, and having *π* = *π*\*k*, lead us to:

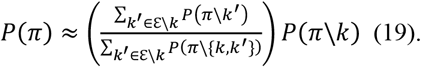

Eq. 19 is obtained conditioned on the insertion of a specific mutation *k* ∈ ε. Thus, it is more accurate to denote *P*(*π*) as *P*^(*k*)^(*π*), and the procedure above must be iterated over every *k* ∈ ε to make the approximation more precise. In other words, Eq. 19 must be iterated *n* times under each of the *n* mutations in ε, which leads to *n* estimated probabilities, averaging of which results in Eq. 10 as desired:

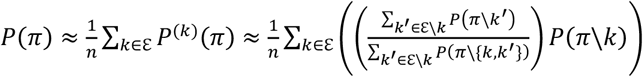

Note that the probability of each pathway in the denominator *P*(*π*\{*k, k*′}) can be calculated by marginalizing over the position of mutation *k* (instead of removing mutation *k* from *π*), which involves summing up the probabilities of *n* − 1 pathways of length *n* − 1 (like what was done in Eq. 12 above). Thus, Eq. 10 remains a function of pathways of length *n* − 1, and the algorithm requires to iterate only one step backward (instead of two steps that Eq. 10 implies).

The pseudocode of the Ensemble R-CBN is presented in **Algorithm 1**, and **Fig. 3** illustrates the procedure for inferring the probability of an example pathway of length *n* = 6. In this example, ε includes 6 mutations, and so the algorithm iterates over 6 different *k* under each of which the pathway probability is calculated and finally their average is considered as the final probability of the given pathway. Under each *k*, the summands of Eq. 10 can be dissected into three separate terms: *i*) the numerator (∑_*k*_′_ε∈\*k*_ *P*(*π*\*k*_′_)), which is obtained by summing up the probabilities of 5 sub-pathways of length 5, each of which lacks one of the 5 remaining mutations in ε\*k, ii*) the denominator (∑_*k*_′_ε∈\*k*_ *P*(*π*\{*k, k*′})), which is obtained by summing up the probabilities of 5 sub-pathways of length 4, each of which lacks the mutation *k* and one of the 5 remaining mutations in ε\*k*, and *iii*) the probability of the sub-pathway of length 5 (*P*(*π*\*k*)), which only lacks the mutation *k*. Note that as mentioned above, the probabilities of the pathways of length *n* − 2 in the denominator of Eq. 10 are quantifiable from the probabilities of pathways of length *n* − 1 by marginalizing over the position of the mutation *k*. For example, the denominators in Fig. 3, for ease of notation, are shown as sum of the probabilities of 5 pathways of length 4, but in practice each denominator is calculated as the sum of the probabilities of 5×5 pathways of length 5, which marginalizes out the position of the mutation *k* (See **Supplementary Fig. S1**).

**Figure 3.**
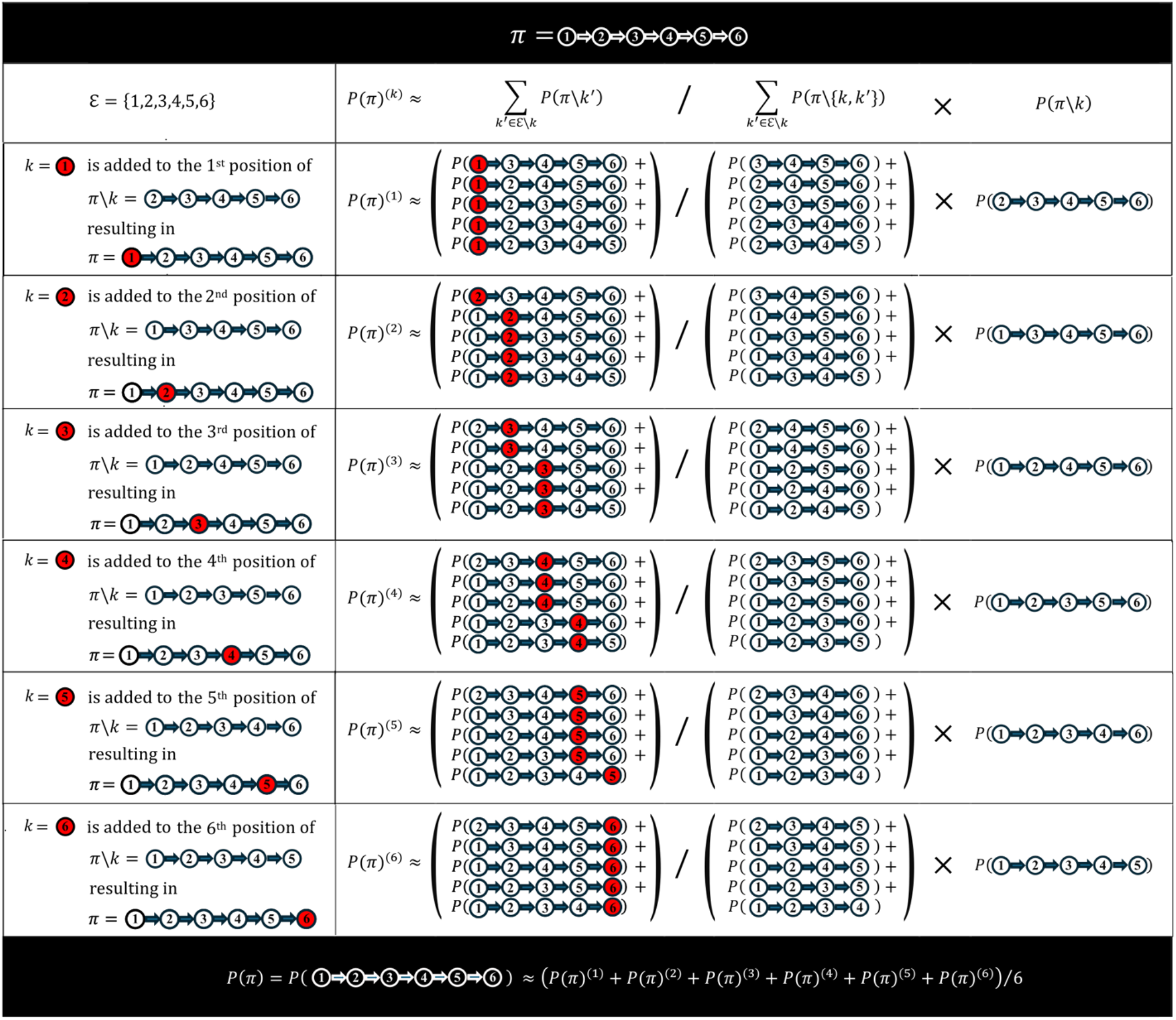
The (Ensemble) R-CBN Algorithm (An Example). A step of the (Ensemble) R-CBN dynamic programming algorithm is illustrated, in which the probability of an example pathway (*π* = 1 → 2 → 3 → 4 → 5 → 6) of length (*n* = 6), which includes 6 mutations from ℰ = {1,2,3,4,5,6}, is derived from the pre-calculated probabilities of pathways of length (*n* = 5). There are six options to arrive at the given pathway *π* of length 6 from one of its sub-pathways of length 5. In each of them, a mutation *k* is inserted at the *k*^*th*^ position along a sub-pathway (*π*\*k*) of length 5, which lacks the mutation *k* (the leftmost column). The pathway probability is approximated using Eq. 10 (the second row), which is illustrated for each of the 6 options (rows 3 to 8). Note that the numerator (∑_*k*_′_∈ℰ\*k*_ *P*(*π*\*k*’)) in this example is obtained by summing up 5 sub-pathways of length 5, each of which lacks one of the 5 mutations in ℰ\*k*. Furthermore, the denominator is obtained by summing up the probability of 5 sub-pathways of length 4 ((∑_*k*′∈ε\*k*_ *P*(*π*\{*k, k*’}))), which are obtained by removing the mutation *k* from the corresponding sub-pathways of length 5 in the numerator. In practice, however, the mutation *k* is not removed from the denominator, but rather its “position” is marginalized out. In other words, each of the five terms in the denominator is a marginal probability obtained by summing up the probabilities of 5 pathways of length 5 in each of which the mutation *k* is inserted into one of the five positions along the corresponding sub-pathway of length 4. Thus, although for ease of notation, the denominator is shown as the sum of the probabilities of 5 pathways of length 4, in practice it is calculated as the sum of the probabilities of 5×5 pathways of length 5 (See **Supplementary Fig. S1**). Finally, this fraction is multiplied by the probability (*P*(*π*\*k*)) of the sub-pathway of length 5 that only lacks the mutation *k*. Thus, the probability of the pathway of length 6 in this example is obtained from the pre-calculated probabilities of pathways of length 5. Similarly, the probability of any given pathway of length *n* can be calculated from the probabilities of pathways of length *n* − 1. This iterative pattern forms the basis for the bottom-up dynamic programming algorithm, which is used in the (Ensemble) R-CBN method.

As explained in **Algorithm 1.1**, the R-CBN-based inference of *N*! Pathways of length *N* is comprised of two parts: Part *I*) (Quartet) R-CBN, which enumerates all potential 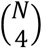 quartets and quantifies all 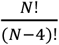 Pathways of length 4 (lines 1-11 in **Algorithm 1.1**). This requires 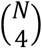 iterations of the (Quartet) R-CBN algorithm, and thus the running time for this part is *O*(*N*^4^).

Part *II*) (Ensemble) R-CBN, which performs *N* − 4 iterative rounds of the bottom-up dynamic programming procedure starting from *n* = 4 until *n* = *N* (Lines 12-29 in **Algorithm 1.1**). For any given pathway of length *n*, the corresponding *n* summands of Eq. 10 are calculated and averaged (lines 22-25 in **Algorithm 1.1**) using the *PathProber* function in **Algorithm 1.2**, which describes how the indices and probabilities of the pathways of length *n* − 1, are found and used to estimate the given pathway probability (line 25). As noted above, these pathways of length *n* − 1 are needed to compose the three distinct elements of Eq. 10, namely: *i)* the numerator (∑_*k*_′_∈G\*k*_ *P*(*π*\*k*′); lines 1-6), *ii)* the denominator (∑_*k*_′_∈G\*k*_ *P*(*π*\{*k, k*′}); lines 7-21), and *iii)* (*P*(*π*\*k*); lines 22-24). The computational task for calculating the probability of a given pathway of length *n* mainly involves finding the index and retrieving the corresponding probabilities of (*n*(*n*(*n* − 1) + 1)) pathways of length *n* − 1 from the previously stored table of pathway probabilities (*Prob*Π′) and pathway IDs (*ID*Π′). Thus, the running time for calculating the probabilities of *N*! pathways of length *N* in part II is *O*(*N*^3^. *N*!). Both parts of the R-CBN algorithm can be efficiently done for pathways of length up to *N* = 10.

#### Algorithm 1.1

Algorithm 1.1 (Ensemble) R-CBN

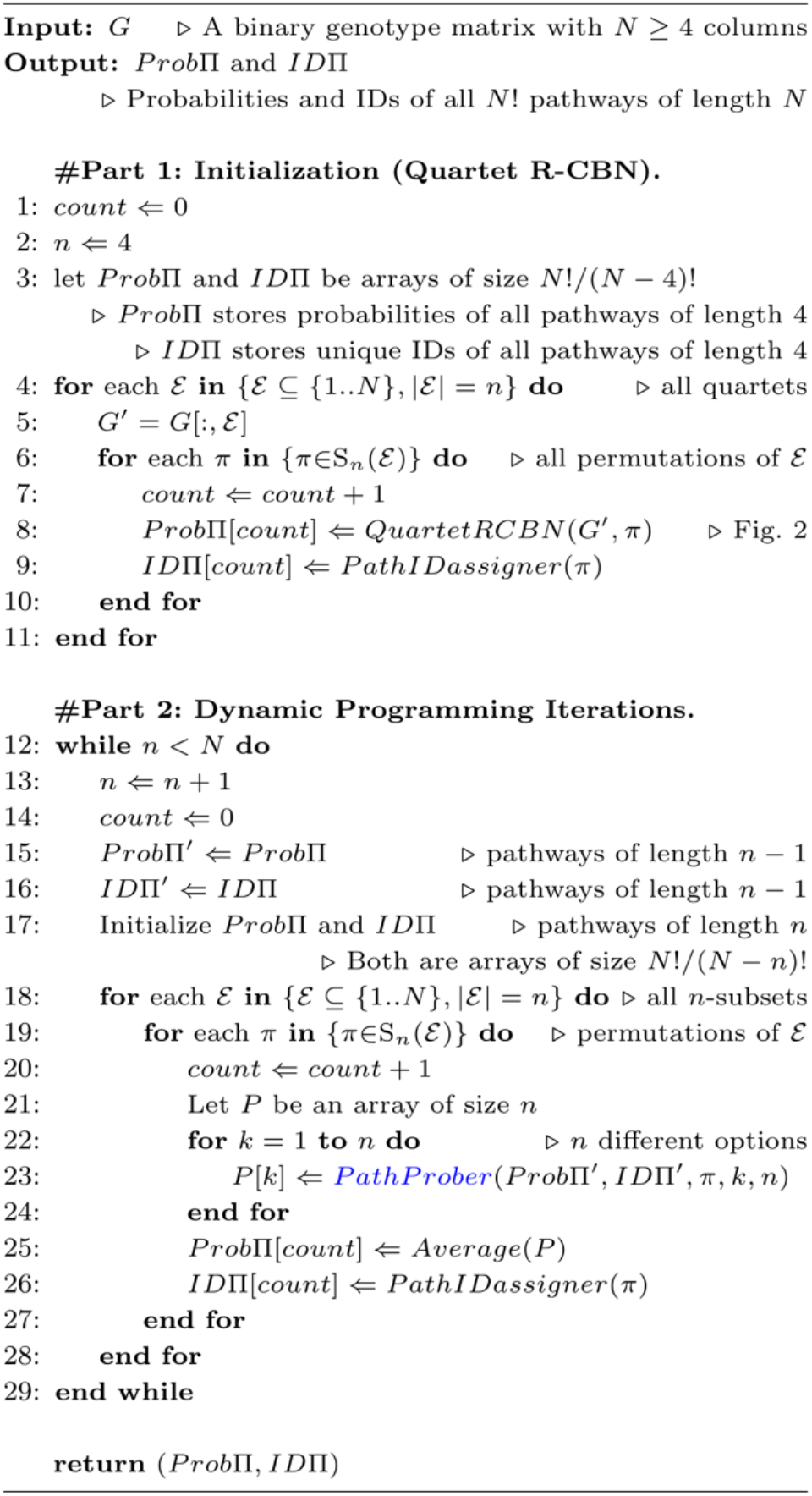

#### Algorithm 1.2

Path Prober

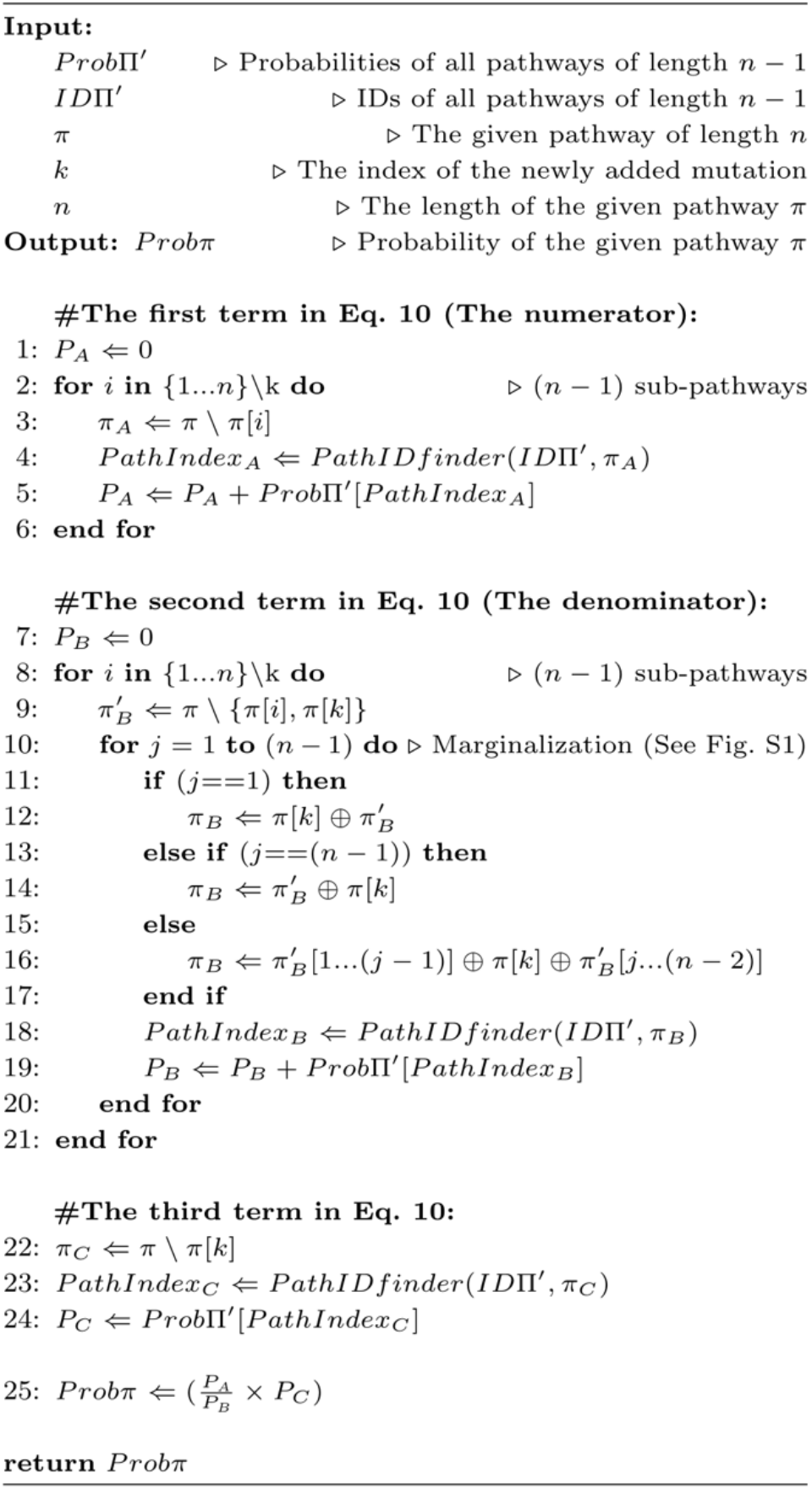

### Evaluation metrics

#### Divergence Score

The Jensen-Shannon Divergence (*JSD*) is used to calculate the divergence between two pathway probability distributions *P*(Π) and *P*′(Π), which is a smoothed version of the Kullback–Leibler divergence:

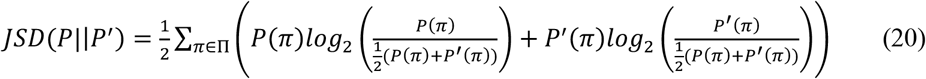

#### Predictability Score

For a given pathway probability distribution *P*(Π) defined over a set of mutational pathways Π, the predictability (Φ_Π_) can be quantified as a function of its entropy (*H*_Π_) as follows [10]:

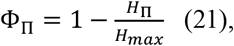

where:

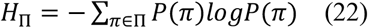

The maximal entropy *H*_*max*_ = *log*(*n*!) is attained when *P*(Π) has uniform distribution, leading to minimal predictability (Φ_Π_ = 0). It is important to note that 0 ≤ Φ_Π_ ≤ 1, and the maximal predictability (Φ_Π_ = 1) or minimal entropy (*H*_Π_ = 0) is attained when only one single pathway among the *n*! potential pathways is feasible.

#### Compatibility Score

Compatibility score (*c*(*π*)) of a given pathway (*π*) is quantified as the fraction of genotypes in the data that are compatible with the mutational ordering specified by the pathway. Recall that the mutational pathway *π* can be represented as an ordered list of *n* + 1 genotypes *g*(*π*) = (*g*_0_, *g*_1_, …, *g*_*n*_), where 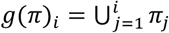. In other words, for any given pathway, there are only *n* + 1 compatible genotypes among the 2^*n*^ potential genotypes. However, the compatibility score differs among pathways, and it depends on the relative frequency (*v*) of the unique set of *n* + 1 compatible genotypes for each pathway. Considering *n* mutations, the pathway-genotype compatibility can be represented as a matrix *M* with *n*! rows and 2^*n*^ columns and each element *M*(*i, j*) of the matrix is 1 if the *j*^*th*^ genotype is compatible with the *i*^*th*^ pathway, and zero otherwise. Having the genotype frequency vector *v* determined from a given cross-sectional mutational data, the compatibility vector (*c*) for the pathways in Π is computed as *c* = *Mv*. Finally, the reliability of a given pathway probability distribution (*P*(Π)) is assessed by computing the Spearman’s rank correlation coefficient (*ρ*) between *c*(Π) and *P*(Π).

### Synthetic Data

To benchmark the CBN models, first I generated synthetic genotypic data under the constraints imposed by pre-specified posets depicted as DAGs of restrictions between mutations. I thus identified 219 unique transitively-closed DAGs of restrictions for a quartet of mutations (*n* = 4), and under each DAG, I generated 200 binary genotypes of length *n* = 4 as follows: First, for each DAG, a genotype set *G* including the subset of genotypes in *U* = {0,1}^4^, which are compatible with the restrictions defined in the given DAG, was determined. Then, 200 genotypes from *G* were sampled deterministically based on the number of mutations contained in each unique genotype in *G*. More precisely, the number of copies of a given genotype in the genotype sample is calculated as follows:

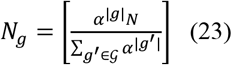

if *g* ∈ *G* and zero otherwise, where |*g*| is the number of mutations in *g*, and I kept *α* = 0.8 and *N* = 200. This sampling strategy ensures that the higher the number of mutations in a genotype, the lower its frequency would be in the sample, which reflects the fact that longer waiting time is required for the arrival of further mutations in the underlying mutation accumulation process.

Furthermore, I repeated this process to generate another set of 219 samples to assess how the CBN models perform in the presence of mutual exclusivity patterns in the genotypic data. Specifically, under each of the 219 DAGs, I determined a genotype set *G*′ ⊂ *G* by randomly choosing a pair of mutations and then excluded all the genotypes in the original set *G* for which both mutations are present. Finally, I repeated the above-mentioned sampling strategy on the updated genotype set *G*′ to generate the second set of samples that contains patterns of mutual exclusivity.

### Simulated Data

To further benchmark the CBN models, I leveraged the data of a previous study [19], which has simulated 100 fitness landscapes representing a pre-specified DAG of restrictions between 7 mutations. In each fitness landscape, a total of 2^7^ = 128 distinct binary genotypes are included, and their fitness is determined based on the restrictions imposed by the DAG and the fitness effects (selection coefficient) of each individual mutation. If according to the given DAG, a genotype is not accessible, its fitness will be set as zero; otherwise, its fitness will be determined based on the set of mutations that it contains. Fitness of the wild-type genotype is always 1, and the fitness of accessible genotypes with a single mutation is 1 + *s*, where *s* is the fitness effect of the mutation and is chosen from a uniform distribution between 0.1 and 0.7. The fitness of accessible genotypes with multiple mutations is 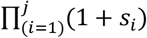, where *s*_*i*_ is the fitness effect of the *i*^*th*^ mutation and *j* is the total number of mutated genes in the given genotype.

While the first approach for inferring the pathway probability distributions can be applied directly to each fitness landscape under the SSWM assumption, the second approach, which is the CBN-based one, operates indirectly on the genotypic data that has been generated under a given fitness landscape. For this purpose, I reused the genotypic data that were previously generated [19] from the fitness landscapes described above through evolutionary simulations using a model [26] implemented in the OncoSimulR package [27]. In each fitness landscape, under different mutation rates (high: 10^−5^ versus low: 10^−6^) and detection regimes (slow versus fast), 20,000 genotypes have been simulated [19], from which I randomly chose 200 genotypes.

### Real data

Finally, I re-used the cancer genomic data that we previously obtained [10] from TCGA [28], which includes samples from primary tumors of untreated patients that we gained access through the cBioPortal platform [29]. This dataset covers 15 distinct cancer types and the number of samples per cancer type varies from 186 to 836. The binary genotype for a given tumor is determined based on the presence (1) or absence (0) of mutations in the 10 most frequently mutated driver genes predicted separately for each cancer type by Mutsig2CV v3.1 [30]. For each of the 15 cancer types, I varied pathway length from *n* = 4 to *n* = 10. For any given *n*, all 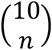 distinct subsets of *n* driver genes were examined, and for each of these subsets the probabilities of *n*! pathways of length *n* were inferred.

## Results

### Benchmarking the CBN models (Synthetic Data)

To systematically benchmark the CBN models, I first focused on pathways of length *n* = 4 for which exhaustive exploration of all transitively closed DAGs of restrictions is feasible. For a gene quartet, the total number of unique transitively closed DAGs is 219, and thus I generated 219 synthetic genotypic samples each corresponding to one of these unique DAGs of length 4 (See Methods). This comprehensive exploration is essential as different DAGs with varied number of edges are required to represent different degrees of evolutionary constraints and hence to cover a wide range of predictability scores. For each synthetic sample, I calculated the CBN-based predictability scores (Φ_CBN_) under each of the four CBN models. Note that the Φ_CBN_ is a function of a pathway probability distribution, which can be used as an indirect quantity to assess the reliability of the pathway probability estimates (See Eq. 21 in Methods). In this analysis, I kept the synthetic genotypic samples error-free, and so they are perfectly compatible with the constraints imposed by their corresponding DAGs, providing an ideal setting to quantify how close the approximate estimations under R-CBN and B-CBN are to their maximum-likelihood-based counterparts, namely the CT-CBN and H-CBN, which are consistent estimators. I thus used CT-CBN as the ground truth to evaluate how precisely the R-CBN and B-CBN estimate the predictability scores. As shown in **Fig. 4a-b**, the predictability scores are the same in H-CBN and CT-CBN, but a slight deviation from the ground-truth is observed in R-CBN and B-CBN methods (**Fig. 4c-d** with Pearson’s R of 0.99 and 0.93, respectively). This deviation is expected as robust frameworks are intended to slightly sacrifice accuracy under ideal conditions for higher robustness under non-ideal conditions characterized by genotypic errors in the data and or violation of the underlying model assumptions.

**Figure 4.**
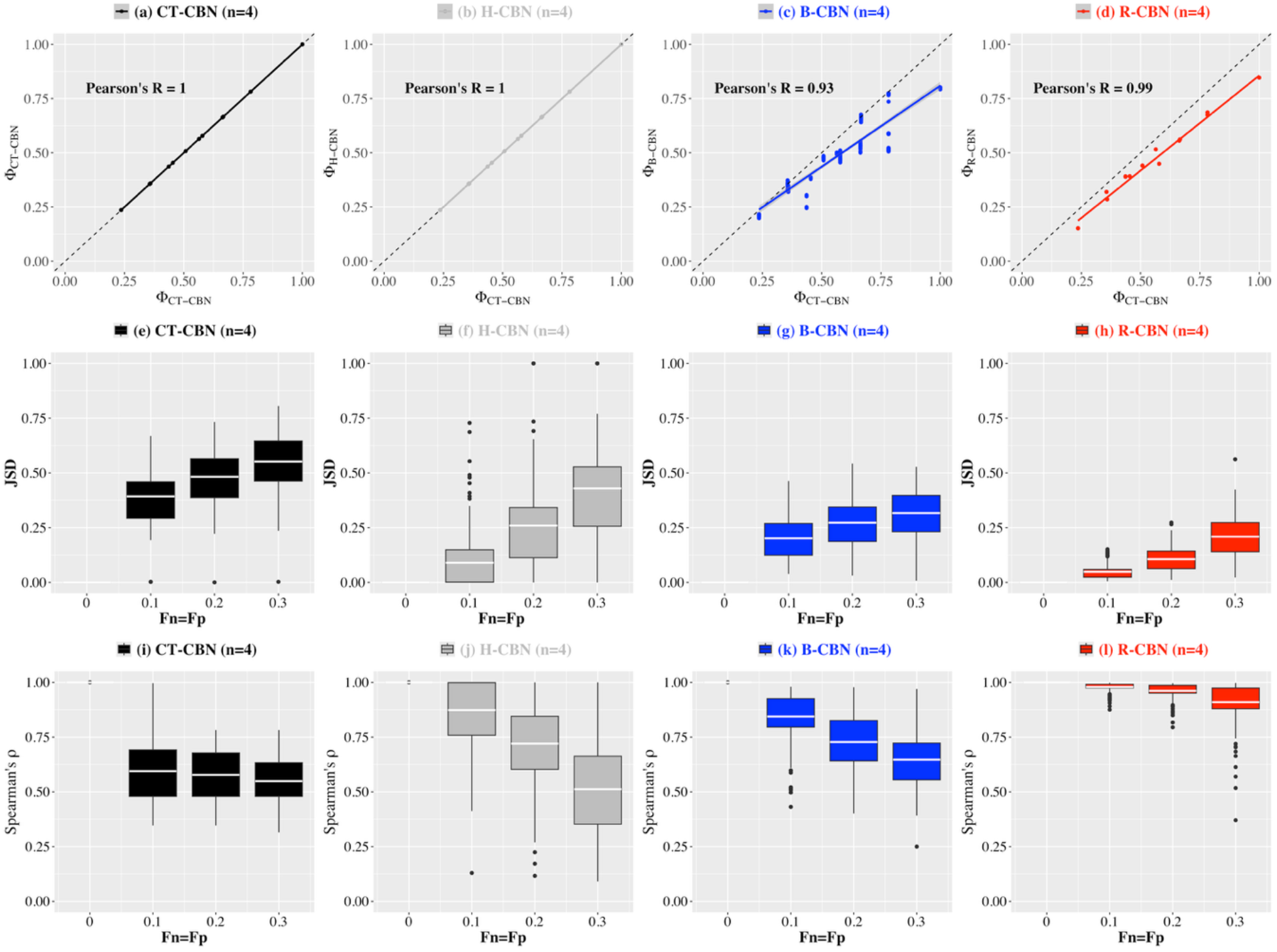
Benchmarking the CBN models (Synthetic Data; *n* = 4). In the upper panels, the horizontal axes indicate the predictability (Φ) quantified using CT-CBN model as the ground truth, and the vertical axes indicate the predictability (Φ) quantified under a) CT-CBN, b) H-CBN, c) B-CBN and d) R-CBN in error-free conditions using 219 synthetic genotypic samples. The box plots show the distribution of the Jensen-Shannon Divergence (*JSD*; the middle panels) and the Spearman’s correlation coefficient (*ρ*; the lower panels) between the pathway probability distributions before and after introducing genotypic errors of a given false positive (*Fp*) and false negative (*Fn*) rate (horizontal axes) using CT-CBN (panels *e* and *i*), H-CBN (panels *f* and *j*), B-CBN (panels *g* and *k*), and R-CBN (*h* and *l*). Note that these analyses are based on pathways of length *n* = 4.

To simulate such non-ideal conditions, I subjected the original synthetic samples to both false-positive (*F*_*p*_; random genotypic changes from 0 to 1) and false negative errors (*F*_*n*_; random genotypic changes from 1 to 0). I repeated this process with different error rates *Fp* = *Fn* ={0.1, 0.2, 0.3} and under each CBN model and for each of the 219 samples, I quantified the Jensen-Shannon Divergence (*JSD*) between the pathway probability distributions inferred before (*P*(Π)) and after (*P*′(Π)) introducing genotypic errors of a given rate (See Eq. 20 in Methods). **Fig. 4e-h** clearly attests to the remarkable robustness of the R-CBN method as it reveals that the JDS values under the R-CBN method remain substantially lower than the other three methods. Under CT-CBN, the JDS values quickly increase by raising error rate; the mean JDS values is already 0.39 at *Fp*(*Fn*) = 0.1 and reaches 0.55 at *Fp*(*Fn*) = 0.3 (**Fig. 4e**). Under H-CBN, the increase in JDS values starts slowly but gradually the rate of increase gets accelerated at higher error rates. For example, the mean JDS at *Fp*(*Fn*) = 0.1 is 0.09 but it reaches 0.43 at *Fp*(*Fn*) = 0.3 (**Fig. 4f**). In contrast, under B-CBN, the JDS values quickly start to increase, and its mean reaches 0.20 at *Fp*(*Fn*) = 0.1, but its rate of increase gradually slows down afterwards such that at *Fp*(*Fn*) = 0.3 its mean JDS is 0.31 (**Fig. 4g**), which is considerably smaller than that of CT-CBN and H-CBN. Under R-CBN, the mean JDS is as low as 0.05 at *Fp*(*Fn*) = 0.1 and its rate of increase remains consistently slow, such that its mean JDS at *Fp*(*Fn*) = 0.3 barely exceeds 0.20 (**Fig. 4h**), which is substantially lower than that of the other methods. Thus far, in this analysis, I have kept *Fp* = *Fn*, but I also examined different combinations of *Fp* and *Fn* including the cases, in which *Fp* ≠ *Fn*, and the results illustrated in **Supplementary Fig. S2** indicate that by increasing error rates either along *Fp* or *Fn*, the above patterns would remain the same for CT-CBN, H-CBN and R-CBN. However, under B-CBN at some combinations of *Fp* and *Fn*, non-monotonic behavior is observed. For example, under B-CBN at *Fp* =0.1 the JDS values at *Fn* =0.2 or *Fn* =0.3 are unexpectedly smaller than those at *Fn* =0.0 or *Fn* =0.1 (**Supplementary Fig. S2j**). Moreover, the JDS values at *Fp* =0.3 are unexpectedly smaller than those at *Fp* =0.2 (**Supplementary Figs. S2k** and **S2l**).

As a complementary analysis, I also evaluated the robustness of the ranking of the pathway probabilities to genotypic errors. For this purpose, under each CBN model, and for each of the 219 samples, I quantified the Spearman’s rank correlation coefficient (*ρ*) between the pathway probability distributions inferred before (*P*(Π)) and after (*P*′(Π)) introducing genotypic errors of a given rate. **Fig. 4i-l** reveals an evident superiority of the R-CBN method in terms of the robustness of the pathway rankings to genotypic errors. Under CT-CBN, the mean *ρ* quickly goes down to 0.59 at *Fp*(*Fn*) =0.1 and remains almost the same at higher error rates (**Fig. 4i**). The mean *ρ* at *Fp*(*Fn*) =0.1 is still as high as 0.87 under H-CBN and 0.84 under B-CBN, but it goes down steeply under H-CBN and relatively slowly under B-CBN, such that at *Fp*(*Fn*) =0.3, the mean *ρ* will be as low as 0.43 under H-CBN and 0.64 under B-CBN (**Fig. 4j-k**). In contrast, under R-CBN, the mean *ρ* at *Fp*(*Fn*) = 0.1 is 0.98, and it remains as high as 0.96 and 0.90 at error rates of 0.2 and 0.3 respectively (**Fig. 4l**). Moreover, the distribution of the *ρ* values under R-CBN is strikingly less spread out as compared to the other methods, which further strengthens evidence in favor of the robustness of R-CBN. Moreover, I examined all combinations of *Fp* and *Fn* including the cases where *Fp* ≠ *Fn*, and the results indicate that the above patterns along both *Fp* and *Fn* would remain the same under the CT-CBN, H-CBN and R-CBN methods, but under B-CBN at some combinations of *Fp* and *Fn*, non-monotonic patterns emerge (**Supplementary Fig. S3**).

Finally, I repeated the above analyses on the second set of synthetic data in which patterns of mutual exclusivity are introduced during genotype generation. The results of the robustness analyses illustrated in **Supplementary Figs. S4, S5 and S6** remain the same as in the first set of genotypes indicating that the presence of mutual exclusivity does not have a noticeable effect on the robustness of the inferred pathway probability distributions to genotypic errors. However, under ideal conditions, the deviation from the ground truth in terms of the predictability scores is more pronounced in the presence of mutual exclusivity, especially under the B-CBN method (**Supplementary Figs. S4c**).

### Benchmarking the CBN models (Simulated Data)

Next, I focused on genotype data which are sampled through evolutionary simulations under 100 fitness landscapes each representing a given DAG of restrictions among *n* = 7 mutations (see Methods). This dataset provides a biologically more realistic ground truth to benchmark the CBN models with, and this analysis is feasible particularly because of the fact that the pathway probability distributions can be quantified both *i*) using an evolutionary model under SSWM assumption (Eq. 4 in Methods), and *ii*) using a CBN-based model (Eq. 6 in Methods). In this setting, performance of the CBN models can be evaluated based on how closely they can reflect the underlying fitness landscape (**Fig. 1**). Thus, I quantified predictability using both the fitness-landscape-based SSWM model (Φ_SSWM_) and the CBN-based model (Φ_CBN_) for each of the 100 fitness landscapes under each of the four CBN models. First, I evaluated the CBN models using pathways of length *n* = 4, which required considering 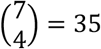 gene quartets and averaging the corresponding 35 Φ_SSWM_ and 35 Φ_CBN_ scores for each fitness landscape. Thus, under each CBN model, this analysis resulted in two vectors Φ_SSWM_ and Φ_CBN_ of length 100 each element of which corresponds to one of the 100 fitness landscapes. **Fig. 5a** shows a nearly perfect correlation between Φ_SSWM_ and Φ_CBN_ under the CT-CBN model (with Pearson’s *R* = 0.97), and relatively strong correlations (with Pearson’s R near 0.9) under H-CBN, B-CBN and R-CBN as well (**Fig. 5b-d**), which confirms that all CBN models can accurately reflect the underlying fitness landscapes in ideal conditions (error-free genotypic data) for pathways of length *n* = 4.

**Figure 5.**
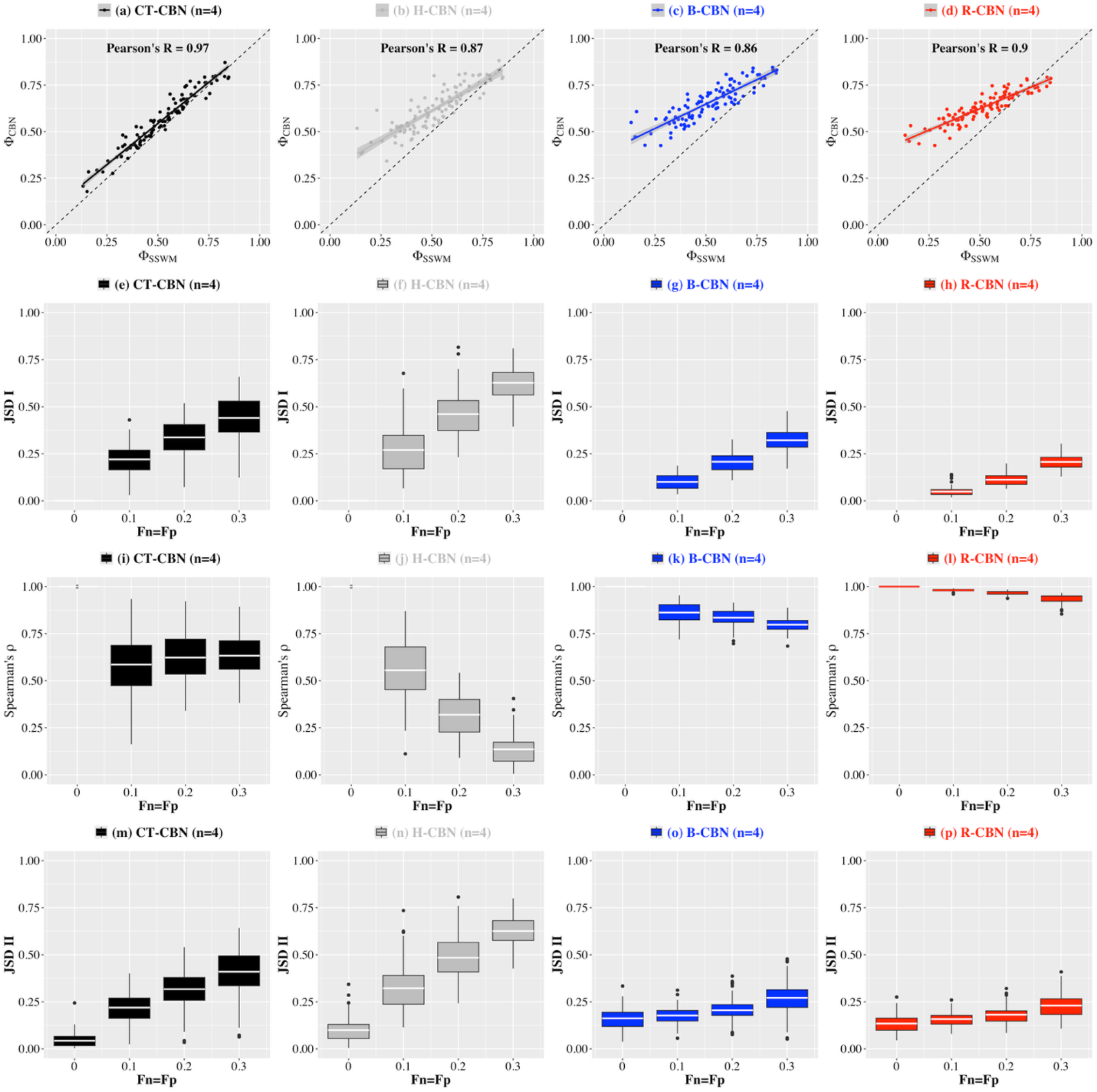
Benchmarking the CBN models (Simulated Data; *n* = 4). In the panels of the first row, the horizontal axes indicate the predictability scores (Φ_SSWM_) obtained using the pathway probability distributions inferred from 100 representable fitness landscape under the evolutionary SSWM-based model, while the vertical axes indicate the predictability scores (Φ_CBN_) obtained under a) CT-CBN, b) H-CBN, c) B-CBN and d) R-CBN models, which have been trained on the genotypic data resulting from evolutionary simulations with low mutation rates (10^−6^) and slow detection regimes (see Methods).The box plots show the distribution of the Jensen-Shannon Divergence (*JSD I*; the second row) and the Spearman’s correlation coefficient (*ρ*; the third row) between the pathway probability distributions before and after introducing genotypic errors of a given false positive (*Fp*) and false negative (*Fn*) rate (horizontal axes) using CT-CBN (panels *e* and *i*), H-CBN (panels *f* and *j*), B-CBN (panels *g* and *k*), and R-CBN (*h* and *l*). The box plots in the fourth row show the distribution of the Jensen-Shannon Divergence (*JSD II*) between the pathway probability distributions calculated using the SSWM evolutionary model and their corresponding CBN model (panels *m* to *p*). Note that this analysis is based on pathways of length *n* = 4, and so for each fitness landscape 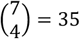 quartets are considered and the final metrics are averaged among them.

I also benchmarked the robustness of the CBN models to genotypic errors. Thus, I subjected the genotypes to false positive (*Fp*) and false negative (*Fn*) errors of a given rate, and under each CBN model and for each of the 100 samples, I quantified the *i)* Jensen-Shannon Divergence (*JSD I*) and *ii)* Spearman’s rank correlation coefficient (*ρ*) between the pathway probability distributions inferred before (*P*(Π)) and after (*P*′(Π)) introducing genotypic errors. **Fig. 5e-l** reaffirms the remarkable robustness of the (Quartet) R-CBN algorithm to genotypic errors. Under CT-CBN and H-CBN, the *JSD I* quickly grows by raising error rates (**Fig. 5e-f**), but under B-CBN and R-CB it stays low and grows slowly (**Fig. 5g-h**). Furthermore, by increasing error rates, the Spearman’s *ρ* is compromised under CT-CBN (**Fig. 5i**) and more severely under H-CBN (**Fig. 5j**), while it remains strong under B-CBN (**Fig. 5k**) and R-CBN (**Fig. 5l**). For example, at *Fp*(*Fn*) = 0.3 the mean *ρ* under R-CBN and B-CBN stays as high as 0.93 and 0.80 respectively, while under CT-CBN and H-CBN it goes down to 0.63 and 0.13 respectively. Moreover, the strikingly lower variance of both the *JSD I* and *ρ* values under R-CBN and B-CBN as compared to those under CT-CBN and H-CBN further attests to their remarkable robustness. Additionally, I calculated the Jensen-Shannon Divergence (*JSD II*) between the pathway probability distributions inferred under the SSWM-based evolutionary model (*P*(*Π*_*f*_)) and under any of the four CBN models (*P*(Π_≤_)). As **Fig. 5m-p** indicates, the *JSD II* values at ideal conditions (*Fp*(*Fn*) = 0) are slightly higher under B-CBN and R-CBN as compared to those under CT-CBN and H-CBN. However, as error rates increase, the *JSD II* values steeply grow under CT-CBN and H-CBN, while they stay low and grow slowly under R-CBN and B-CBN, which provide further evidence in favor of their robustness. Note that in this analysis so far, the genotype samples were obtained through evolutionary simulations under low mutation rate (10^-6^) and slow detection regimes. I repeated the above analysis using samples obtained under high mutation rate (10^-5^) and fast detection regime, and as **Supplementary Fig. 7** indicates, the above patterns stay the same under this condition. Thus, the evolutionary simulation data analysis corroborates the main conclusions from the synthetic data analysis for pathways of length *n* = 4, which further confirms the validity of the (Quartet) R-CBN method.

To benchmark the CBN models including the (Ensemble) R-CBN, analyzing longer pathways (*n* > 4) is needed, and because the fitness landscapes are defined for *n* = 7 genes, the simulated data enables this goal. The most direct way to assess the accuracy of the (Ensemble) R-CBN approximate approach is to compare it with its own exact version. As mentioned in the Methods section, the exact inference under R-CBN requires exhaustive examination of all potential posets of a given length *n*, which quickly becomes infeasible at *n* ≥ 6. Nevertheless, exact R-CBN inference is still feasible at *n* = 5 (Quintet R-CBN) albeit computationally more demanding than at *n* = 4 (Quartet R-CBN). Therefore, the accuracy of one step of the dynamic programming algorithm in (Ensemble R-CBN) can be directly assessed at *n* = 5 by comparing the pathway probability distributions inferred under the exact (Quintet) R-CBN with those inferred under the approximate Ensemble R-CBN using both Jensen-Shannon Divergence (*JDS*) and Spearman’s rank correlation coefficient (*ρ*). For this purpose, I considered all 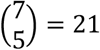 quintets per each of the 100 fitness landscapes, and so calculated 21 × 100 *JDS* and *ρ* values separately under two distinct evolutionary scenarios and four different error rates. The results are summarized in **Table. 1**, which indicates that under all conditions, the *JDS* values stay close to 0, and the *ρ* values remain almost 1, and so the difference between the two inferential schemes is negligible, which validates the accuracy of the approximate (Ensemble) R-CBN at *n* = 5.

**Table 1.** Exact vs. Approximated R-CBN (*n* = *5*). The mean and standard deviation of the Jensen-Shannon Divergence (*JSD*; the second and fourth columns) and the Spearman’s rank correlation coefficient (*ρ*; the third and fifth columns) between the exact-RCBN-based versus the approximated R-CB-based pathway probability distributions are listed. These statistics are based on 100 simulated genotypic data, each of which is derived from one of the 100 representable fitness landscapes under *i*) low mutation rate (10^−6^) and slow detection regimes (the second and third columns), and *ii*) high mutation rates (10^−5^) and fast detection regimes (the fourth and fifth columns), after being subjected to various false positive (*Fp*) and false negative (*Fn*) rates (the first column). Note that this analysis is based on pathways of length *n* = 5, and so 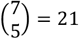 quintets are considered per fitness landscape.

| Exact R-CBN vs. Approximated R-CBN ($n = 5$) | | | | |
| --- | --- | --- | --- | --- |
| $(Fp, Fn)$ | 100 Representable Fitness Landscapes<br>Low Mutation Rate & Slow Detection Regime | | 100 Representable Fitness Landscapes<br>High Mutation Rate & Fast Detection Regime | |
| | $JSD$<br>(Mean ± SD) | Spearman's $\rho$<br>(Mean ± SD) | $JSD$<br>(Mean ± SD) | Spearman's $\rho$<br>(Mean ± SD) |
| (0.0, 0.0) | **0.05** ± 0.018 | **0.987** ± 0.004 | **0.06** ± 0.024 | **0.986** ± 0.005 |
| (0.1, 0.1) | **0.03** ± 0.012 | **0.991** ± 0.007 | **0.03** ± 0.010 | **0.990** ± 0.007 |
| (0.2, 0.2) | **0.02** ± 0.006 | **0.993** ± 0.003 | **0.03** ± 0.006 | **0.993** ± 0.003 |
| (0.3, 0.3) | **0.02** ± 0.008 | **0.992** ± 0.009 | **0.02** ± 0.008 | **0.993** ± 0.005 |

Furthermore, after three iterations of the (Ensemble) R-CBN algorithm, I calculated the probability distributions for pathways of length *n* = 7 and compared them with those obtained under the other CBN models by measuring the Φ_SSWM_, Φ_CBN_, *JSD I, JSD II* and Spearman’s *ρ* for each of the 100 fitness landscapes under two evolutionary scenarios and four different error rates. **Fig. 6** indicates that the patterns described above for (Quartet) R-CBN at *n* = 4 remains the same for (Ensemble) R-CBN at *n* = 7. However, comparing **Figs 5** and **6** reveals some noticeable differences under some of the other CBN methods. While the correlation between Φ_SSWM_ and Φ_CBN_ at *n* = 7 remains strong under CT-CBN, H-CBN and R-CBN, it is considerably compromised under B-CBN (**Fig. 6a-d**). Moreover, at *n* = 7 the *JSD I* values under CT-CBN, H-CBN and B-CBN are considerably higher than those at *n* = 4, and so at *n* = 7, the robustness of B-CBN is no longer comparable to that of R-CBN method (**Fig. 6e-h**). Similarly, the Spearman’s *ρ* at *n* = 7 is more severely compromised than at *n* = 4 under CT-CBN and B-CBN, and thus none of the CBN models remain comparable to R-CBN at *n* = 7 in terms of robustness of the pathway rankings to genotypic errors (**Fig. 6i-l**). At *n* = 7 the *JSD II* values are in general higher than those at *n* = 4 under all CBN models, but the main patterns remain the same as in *n* = 4 (**Fig. 6m-p**). In ideal error-free conditions (*Fp*(*Fn*) = 0.0), the *JDS II* values under R-CBN and especially under B-CBN are higher than those under CT-CBN and H-CBN. However, as error rates grow, the *JSD II* values under CT-CBN and H-CBN quickly surpass those under B-CBN and R-CBN. Finally, I repeated these analyses using genotypic data simulated under high mutation rates and fast detection regimes, and as **Supplementary Fig. 8** indicates, the main conclusions stay almost the same. Thus, under various conditions and using several metrics, the results attest to the remarkable robustness of the Ensemble R-CBN method to genotypic errors at pathway length *n* = 7.

### Benchmarking the CBN models (Real Data)

Finally, I analyzed real cancer genomic data from The Cancer Genome Atlas (TCGA), which includes 15 cancer types, each of which contains hundreds of genomic samples from primary tumors of untreated cancers. For each cancer type, the genotypes are determined based on the presence (1) or absence (0) of mutations in the 10 most frequently mutated driver genes. For the real data, the underlying fitness landscapes (as in simulated data) or the underlying DAG of restrictions (as in synthetic data) are not known in advance, so an alternative approach for benchmarking the CBN models is needed. Thus, as a ground truth I quantified the pathway compatibility score (*c*(*π*)) defined as the fraction of genotypes in the data that are compatible with the mutational orders specified by a given pathway (*π*) (see Methods). In general, only *n* + 1 genotypes among the 2^*n*^ potential genotypes are compatible with a given pathway. Two genotypes (the wild-type (all 0) and the fully mutated genotype (all 1)) are generally compatible with every pathway, and there are a unique set of *n* − 1 genotypes specifically compatible with a given pathway. As an example in **Fig 7a**, at *n* = 4, genotypes are defined based on mutations in four genes namely KRAS, TP53, CDKN2A and RREB1, and the compatibility of all 2^4^ = 16 genotypes on each of the 4! = 24 potential pathways of length *n* = 4 are shown as a binary matrix (*M*) each row of which has five 1s corresponding to the genotypes compatible with the given pathway (**Fig. 7a**). The compatibility score (*c*) of a pathway on a given sample of genotypes can be calculated according to the relative frequency (*v*) of its corresponding compatible genotypes (**Fig. 7b**), and for all potential pathways of length *n* denoted as Π = *S*_*n*_, the vector of compatibility scores (*c*(Π)) is calculated as *c* = *Mv* (**Fig. 7c**). For a given pathway *π* with a higher compatibility score (*c*(*π*)), a higher probability (*P*(*π*)) is expected, and so the Spearman’s rank correlation coefficient (*ρ*) between (*c*(Π)) and (*P*(Π)) can be used as a criterion to benchmark the CBN models.

**Figure 6.**
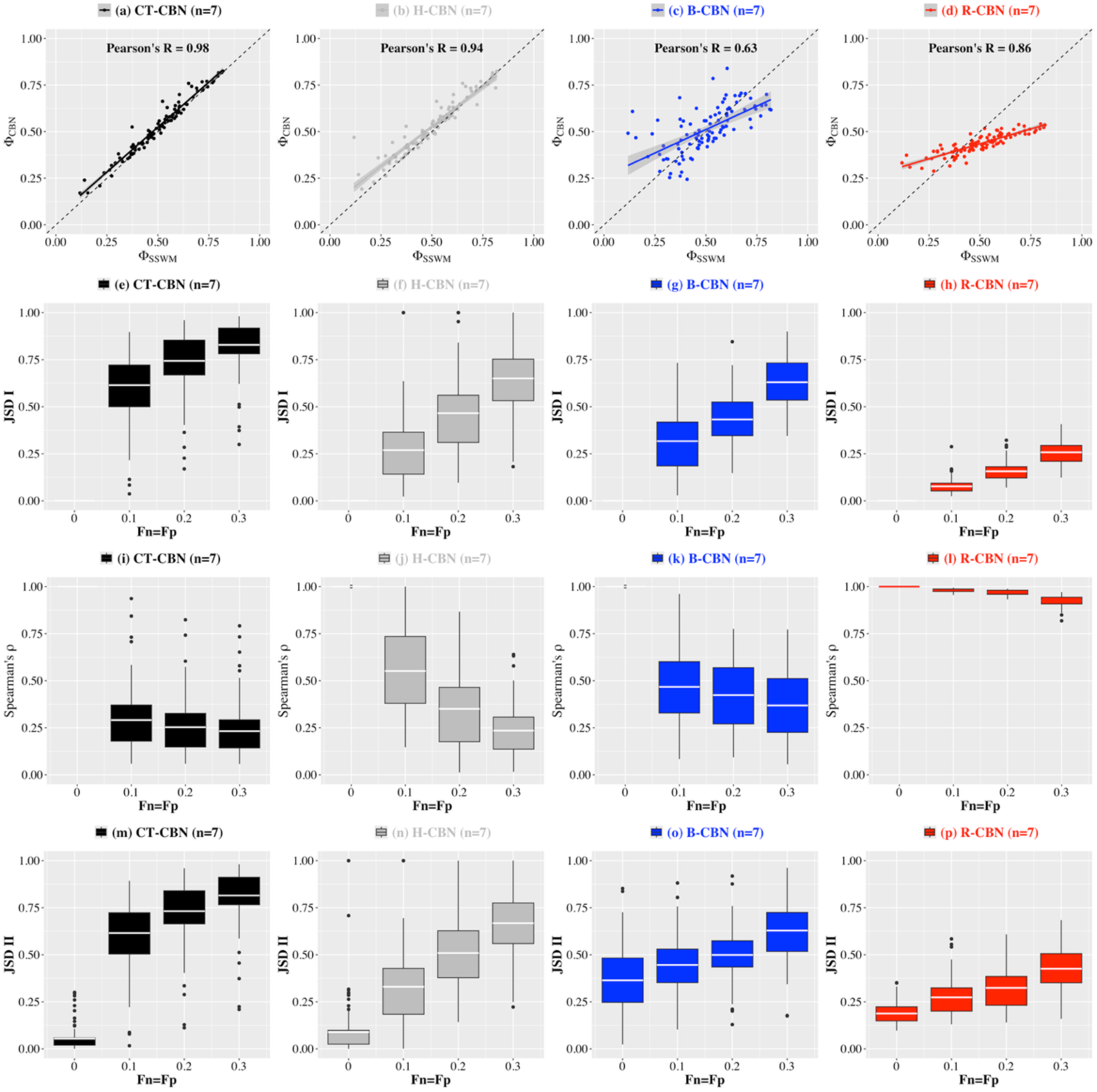
Benchmarking the CBN models (Simulated Data; *n* = 7). In the panels of the first row, the horizontal axes indicate the predictability scores (Φ_SSWM_) obtained using the pathway probability distributions inferred from 100 representable fitness landscape under the evolutionary SSWM-based model, while the vertical axes indicate the predictability scores (Φ_CBN_) obtained under a) CT-CBN, b) H-CBN, c) B-CBN and d) R-CBN models, which have been trained on the genotypic data resulting from evolutionary simulations with low mutation rates (10^−6^) and slow detection regimes (see Methods).The box plots show the distribution of the Jensen-Shannon Divergence (*JSD I*; the second row) and the Spearman’s correlation coefficient (*ρ*; the third row) between the pathway probability distributions before and after introducing genotypic errors of a given false positive (*Fp*) and false negative (*Fn*) rate (horizontal axes) using CT-CBN (panels *e* and *i*), H-CBN (panels *f* and *j*), B-CBN (panels *g* and *k*), and R-CBN (*h* and *l*). The box plots in the fourth row show the distribution of the Jensen-Shannon Divergence (*JSD II*) between the pathway probability distributions calculated using the SSWM evolutionary model and their corresponding CBN model (panels *m* to *p*). Note that this analysis is based on pathways of length *n* = 7.

**Figure 7.**
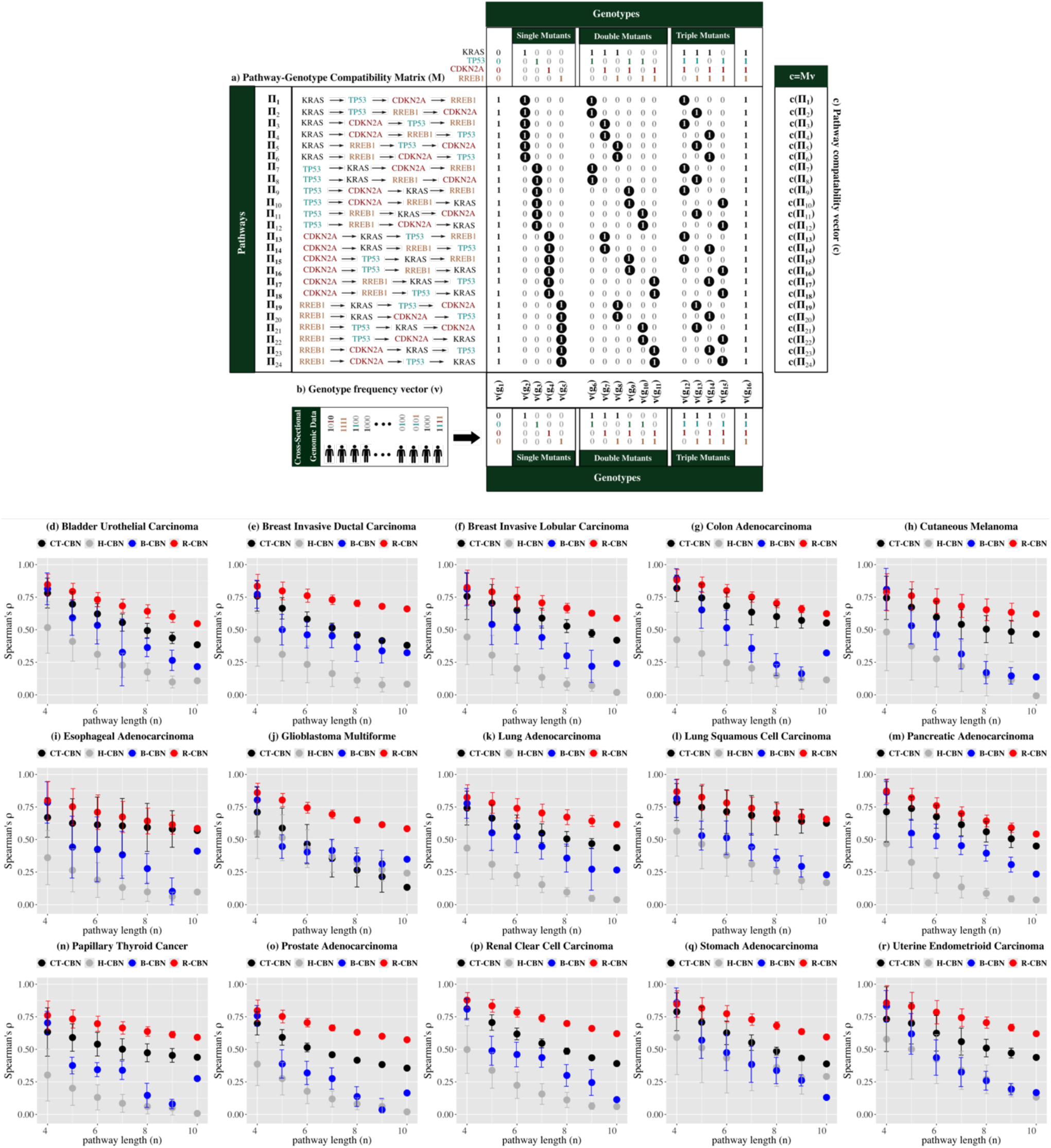
Pathway-Genotype Compatibility Analysis (Real Data; 4 ≤ *n* ≤ 10). (a) An example of a quartet including KRAS, TP53, CDKN2A, and RREB1 genes along with the set of all 2^3^ = 16 binary genotypes and all 4! = 24 mutational pathways of length 4 are shown. The 24 × 16 pathway-genotype matrix *M* encodes the compatibility (1) or incompatibility (0) of a given genotype (columns) on a given pathway (rows). There are only 5 compatible genotypes for any given pathway of length *n* = 4. There are always, 2 genotypes (the all-0s and all-1s), which are compatible with every pathway, but the other 3 genotypes (highlighted by black circles) are pathway specific. (b) The relative frequency (*v*) of each of the 16 genotypes is determined from a given cross-sectional mutational data, and (c) the pathway compatibility vector (*c*) is quantified as *c* = *Mv*. Note that this scheme can be generalized to pathways of any length *n*, for which an *n*! × 2^*n*^ pathway-genotype matrix *M* can be constructed, where each pathway will have *n* + 1 compatible genotypes (the all-0s, the all-1s, and *n* − 1 pathway-specific genotypes). Each of the bottom panels (d to r) corresponds to a specific cancer type, in which the vertical axis indicates the Spearman’s rank correlation coefficient (*ρ*) between the compatibility scores (*c*(Π)) and the corresponding pathway probability distribution (*P*(Π)), which are obtained under CT-CBN (black), H-CBN (gray), B-CBN (blue) and R-CBN (red) models trained on TCGA data. A set of 10 driver genes are considered per cancer type, and pathways of length *n* ranging from *n* = 4 to *n* = 10 are examined (horizontal axes). Note that for a given *n*, there are 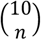 gene subsets resulting in 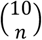 Spearman’s *ρ* values, whose distribution is summarized by its mean (circles) and its standard deviation (error bars).

Having defined the TCGA genotypes based on the presence of mutations in *n* = 10 genes, I was able to systematically vary pathway length from *n* = 4 to *n* = 10, which is required for rigorous assessment of the scalability of the (Ensemble) R-CBN method as compared to the other CBN models. For any given *n*, I examined all 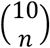 distinct subsets of *n* driver genes, and for each of these subsets, I calculated both the probability (*P*(Π)) and the compatibility (*c*(Π)) of the corresponding *n*! pathways of length *n*, which resulted in 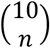 Spearman’s *ρ* values. **Fig. 7d-r** shows the mean and standard deviation of the Spearman’s *ρ* values as a function of *n* separately for each of the 15 cancer types under each of the four CBN models. At *n* = 4, the Spearman’s *ρ* values under the (Quartet) R-CBN are only slightly higher than those of B-CBN and CT-CBN in all cancer types. However, as *n* increases, the superiority of the (Ensemble) R-CBN as compared to the other CBN models becomes more apparent. Under B-CBN, the *ρ* values considerably go down at *n* = 5 and continue diminishing until *n* = 10. Furthermore, under B-CBN occasional fluctuations and high variance of the *ρ* values at higher *n* are observed, which further erode its competitiveness against the (Ensemble) R-CBN. In most cancer types, CT-CBN keeps it’s *ρ* values higher than B-CBN and the rate of decrease of the *ρ* values under CT-CBN is considerably slower than in B-CBN. Nevertheless, the mean *ρ* values under CT-CBN in all cancer types remains below those under R-CBN. For example, at *n* = 7, in 12 out of 15 cancer types (80%), the mean *ρ* values under R-CBN is more than 0.10 higher than that under CT-CBN, and at *n* = 10, in 11 out of 15 cancer types (73.3%), the *ρ* value under R-CBN is more than 0.15 higher than that under CT-CBN. Qualitatively speaking, at larger *n*, only under R-CBN the *ρ* values remain high enough. For example, at *n* = 7, the mean *ρ* values under R-CBN range between 0.66 and 0.75, and so remain moderately high, while under CT-CBN only in 4 out of 15 cancer types (26.6%) the mean *ρ* values at *n* = 7 reach 0.60, and under B-CBN or H-CBN, in none of the cancer types the mean *ρ* values exceeds 0.50. Even at *n* = 10, the *ρ* values under R-CBN remain relatively high as they range between 0.54 and 0.66, while under CT-CBN in only 3 cancer types (20%), the *ρ* value exceeds 0.50. Furthermore, the performance of CT-CBN across cancer types varies dramatically and is not as consistent as R-CBN. For example, in three cancer types, namely Esophageal Adenocarcinoma (**Fig. 7i**), Lung Squamous Cell Carcinoma (**Fig. 7l**) and Pancreatic Adenocarcinoma (**Fig. 7m**), the *ρ* values under CT-CBN are almost as high as those under R-CBN. In contrast, in some cancer types such as Breast Invasive Ductal Carcinoma (**Fig. 7e**) and Glioblastoma Multiforme (**Fig. 7j**), the Δ*ρ* between R-CBN and CT-CBN at *n* = 10 for example, is as large as 0.28 and 0.45 respectively. It is also noticeable that in Glioblastoma Multiforme (**Fig. 7j**), the *ρ* values under CT-CBN stay below than those under B-CBN and even uder H-CBN that has already fallen out of competition in this analysis. Thus, to put it in a nutshell, R-CBN remains the only method that can keep the *ρ* values high enough consistently in all cancer types and at all pathway lengths, which further validates the robustness and scalability of the (Ensemble) R-CBN algorithm.

## Discussion

In this study, I have introduced the R-CBN method for robust and scalable quantification of cancer progression pathways from cross-sectional genomic data. Motivated by the inherent modularity of biological systems [31, 32], first I designed the (Quartet) R-CBN, which aims to infer probabilities of pathways of length *n* = 4 by exhaustively considering all potential posets of size *n* = 4. Quartet R-CBN utilizes a two-tier weighting scheme operating respectively on posets and pathways, which enables close approximation of the predictability scores and exhibits remarkable robustness of pathway probability distributions to genotypic errors. Next, by identifying an iterative pattern, which allowes a bottom-up dynamic programming algorithm (the Ensemble R-CBN), I enabled robust quantification of longer pathways starting from pathways of length *n* = 4. Note that this algorithm operates directly on the pathway probabilities, without the need for exploring the space of potential posets, which becomes progressively huge as poset size *n* increases. This is a remarkable shortcut as compared to the alternative approximation strategies such as the simulated annealing in H-CBN or the MCMC sampling in B-CBN, which rely on exploring the vast space of potential posets.

R-CBN consistently demonstrates superior robustness to genotypic errors on different data types including synthetic data representing a wide range of evolutionary constraints, simulated data covering hundreds of fitness landscapes and real cancer genomic data including several cancer types. Furthermore, the (Ensemble) R-CBN method exhibits reliable inference of probabilities of pathways of various length and hence allows a scalable inferential scheme. The maximum-likelihood based approaches, such as CT-CBN and H-CBN, enable nearly perfect inference under ideal error-free conditions. However, using various metrics in this study, my analyses clearly demonstrate their lack of robustness to genotypic errors. Furthermore, the Bayesian B-CBN approach cannot offer a steadily solid inference, either. For pathways of length *n* = 4, the robustness of B-CBN approach is comparable to that of (Quartet) R-CBN on both synthetic data (**Fig. 4**) and simulated data (**Fig. 5**). However, B-CBN does not seem to be scalable as its robustness to genotypic error gets substantially compromised as pathway length (*n*) increases (**Fig. 6** and **Fig 7**). Thus, R-CBN remains the only method that delivers reliable inference of cancer progression pathways across various conditions. Nevertheless, it is important to note that these observations do not challenge the validity of the previous CBN models. In this study, these methods are basically evaluated in a new domain for which they were not originally designed for. In principle, the former CBN models were intended to generate insights directly from the inferred DAG of restrictions without considering the pathways and their probabilities. In contrast, R-CBN here is an attempt to repurpose the CBN model specifically to excel in this new domain, which is centered on pathway probability distributions and their robustness to genotypic errors.

Importantly, the R-CBN model yields a strikingly robust inference of pathway rankings. This could be especially helpful in navigating into pathway-level biomarkers of cancer progression [33], which could further empower personalized medicine by facilitating mechanistic interpretability and inspiring new evolutionary-principled disease staging strategies [34]. This would warrant follow-up investigations into the repeated evolutionary trajectories [35–37], which would require rigorous benchmarking with single-cell or multi-region samples and longitudinal data. Perhaps more importantly, R-CBN can also potentially be utilized in (causal) predictive modeling by enabling novel pathway-centric disease representation strategies [18]. In this regard, the fact that R-CBN retains the ability to reflect the underlying fitness landscapes directly from cross-sectional genomic data is promising. Establishing a predictive framework that can model how fitness landscapes are reshaped upon perturbation can open new avenues in precision medicine [38], especially in the advent of single-cell genomics, where cross-sectional genomic snapshots are pervasively available [39]. However, achieving this is contingent on proving the ability of the R-CBN model to accurately reflect the underlying perturbation-induced fitness landscapes [40], which calls for further investigations.

Regarding algorithmic efficiency, the running time for pathways of length *n* is *O*(*n*^4^) under the (Quartet) R-CBN, and *O*(*n*^S^. *n*!) under the (Ensemble) R-CBN. For pathways of length up to *n* = 10, the inference is efficient, but both memory and computational issues will start to arise when storing and calculating the probabilities of *n*! pathways at *n* > 10. These computational hurdles are addressable if we take advantage of the sparsity of the pathway probabilities, because only a tiny fraction of the pathways tends to have a probability significantly above zero. However, there is no strong biological motivation for calculating probabilities of pathways of length *n* > 10, especially because epistatic influence on fitness landscapes is dramatically declined at epistatic order higher than *n* = 10 [41], and hence the associated pathway probability distributions at *n* > 10 are not likely to bring any additional information. Furthermore, using alternative levels of abstraction such as sub-processes or sub-systems levels [42], instead of the gene-level one used in the present study, more systematic analyses will become feasible at lower *n*. Finally, it is important to note that CBN-based modeling is not necessarily limited to point mutations, but rather it is extendable to accommodate alternative molecular changes as long as we are able to represent the underlying processes as the stepwise accumulation of binary changes, which are common in molecular biology, such as active (1) versus inactive (0) forms, or phosphorylated (1) versus dephosphorylated (0) states; etc. This opens further opportunities for integrating multiple data modalities, which is particularly essential in the multi-omics era.

In summary, I have introduced the R-CBN algorithm, which enables robust and scalable inference of cancer evolutionary trajectories that will pave the way for the emergence of novel evolutionary-informed modeling strategies in precision cancer medicine.

## Code availability

Codes and data will be available upon publication.

## Acknowledgments

Computational resources for this project were provided by the Texas Advanced Computing Center (TACC).

## Supplementary Figures

**Supplementary Figure S1.**
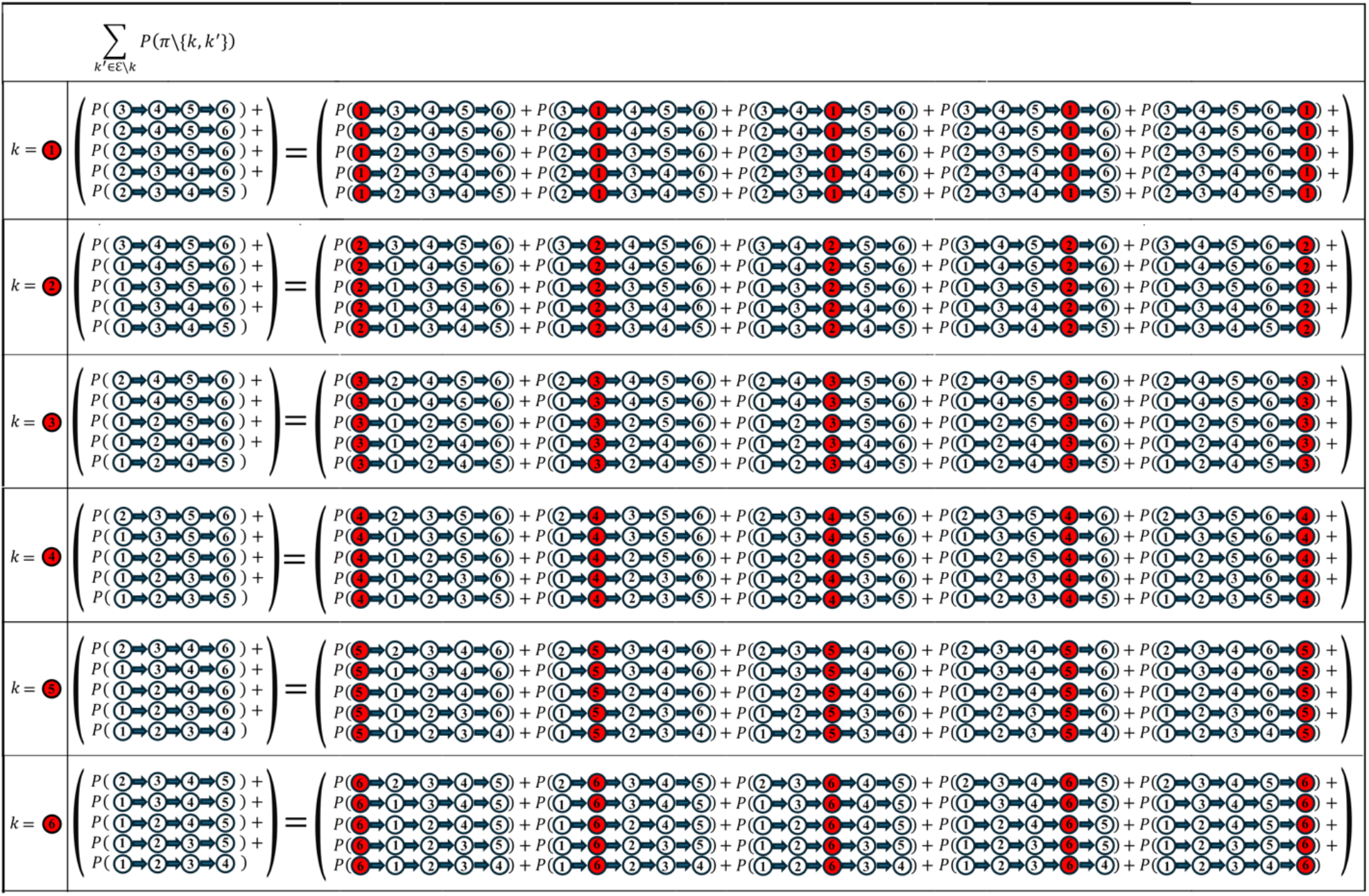
Denominator of Eq. 10 (Marginalization over the position of mutation *k*). In this example, the denominator of Eq. 10 is illustrated as the sum of probabilities that are obtained by marginalizing over the “position” of mutation *k*. In **Fig. 3**, for ease of notation, the denominator is shown as the sum of the probability of sub-pathways of length 4 ((∑_*k*_′_∈ℰ\*k*_ *P*(*π*\{*k, k*′}))). In practice, however, the mutation *k* is not removed from the denominator, but rather its “position” is marginalized out. In other words, each of the five terms in the denominator is a marginal probability obtained by summing up the probabilities of 5 pathways of length 5 in each of which the mutation *k* is inserted into one of the five positions along the corresponding pathway of length 4. Thus, the denominator is calculated as the sum of the probabilities of 5×5 pathways of length 5. Note that in this figure six denominators corresponding to each of the six *k* ∈ {1,2,3,4,5,6} are illustrated, which are the equivalents of the example provided in **Fig. 3**.

**Supplementary Figure S2.**
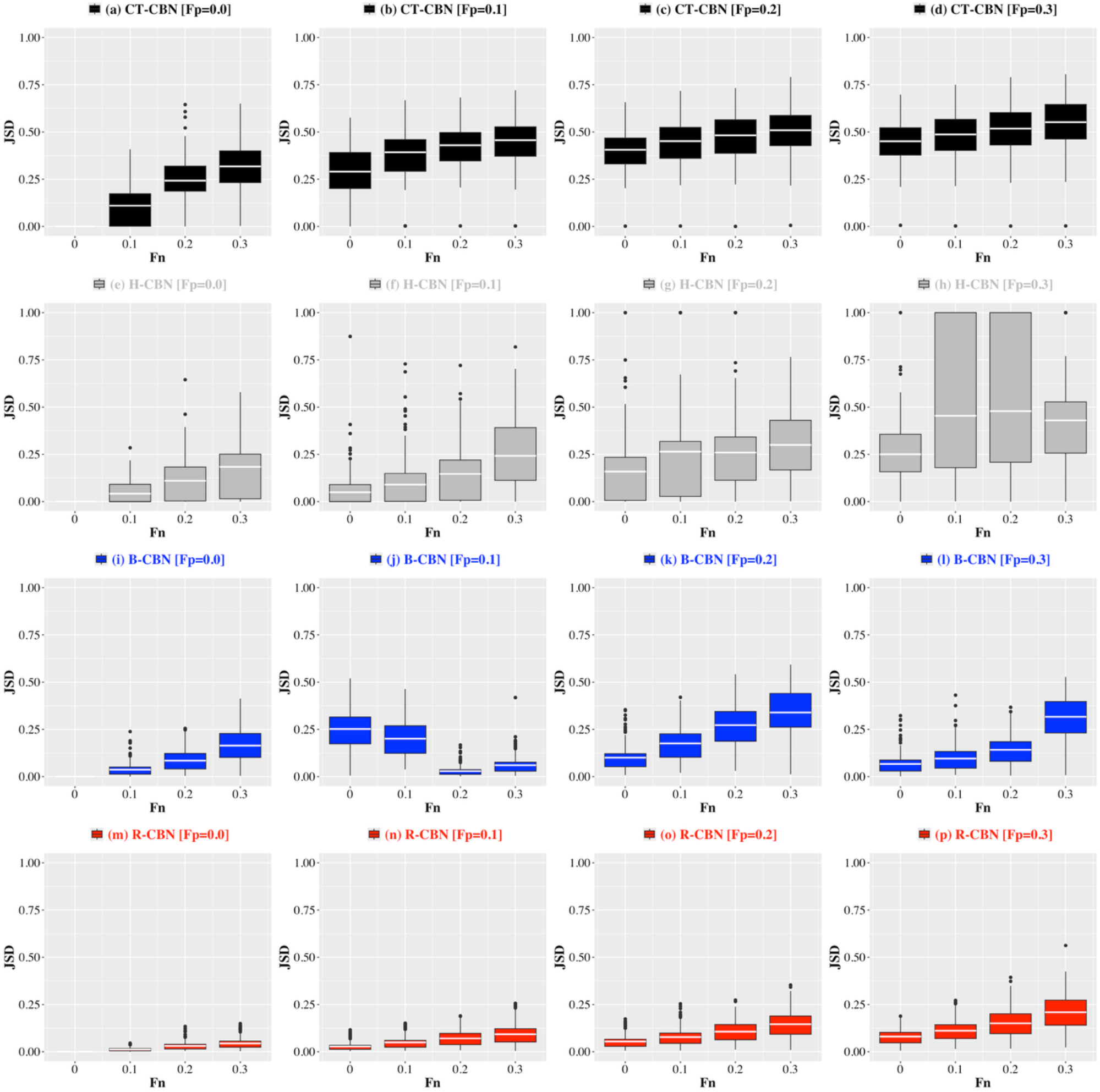
Robustness analysis I (Synthetic Data, *n* = 4 [All combinations of *Fn* and *Fp*]). The box plots show the distribution of the Jensen-Shannon Divergence (*JSD*) between the pathway probability distributions before and after introducing genotypic errors of a given false positive (*Fp*) rate (specified in the title of each panel) and false negative (*Fn*) rate (horizontal axes) using *i)* CT-CBN (black, the first row), *ii)* H-CBN (gray, the second row), *iii)* B-CBN (blue, the third row), and *iv)* R-CBN (red, the fourth row). Note that the synthetic data analyses are based on pathways of length *n* = 4.

**Supplementary Figure S3.**
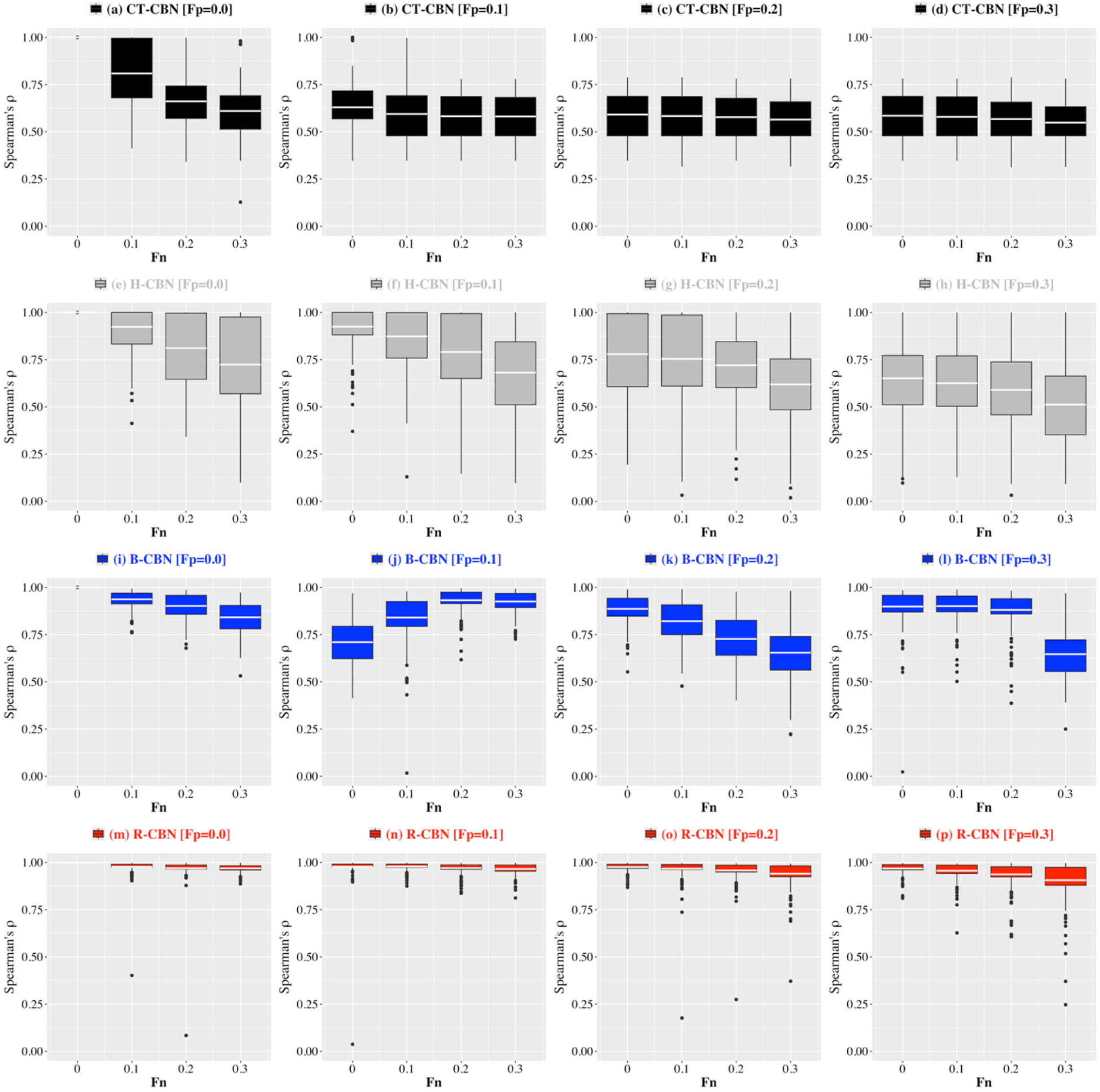
Robustness analysis II (Synthetic Data, *n* = 4 [All combinations of *Fn* and *Fp*]). The box plots show the distribution of the Spearman’s correlation coefficient (*ρ*) between the pathway probability distributions before and after introducing genotypic errors of a given false positive (*Fp*) rate (specified in the title of each panel) and false negative (*Fn*) rate (horizontal axes) using *i)* CT-CBN (black, the first row), *ii)* H-CBN (gray, the second row), *iii)* B-CBN (blue, the third row), and *iv)* R-CBN (red, the fourth row). Note that the synthetic data analyses are based on pathways of length *n* = 4.

**Supplementary Figure S4.**
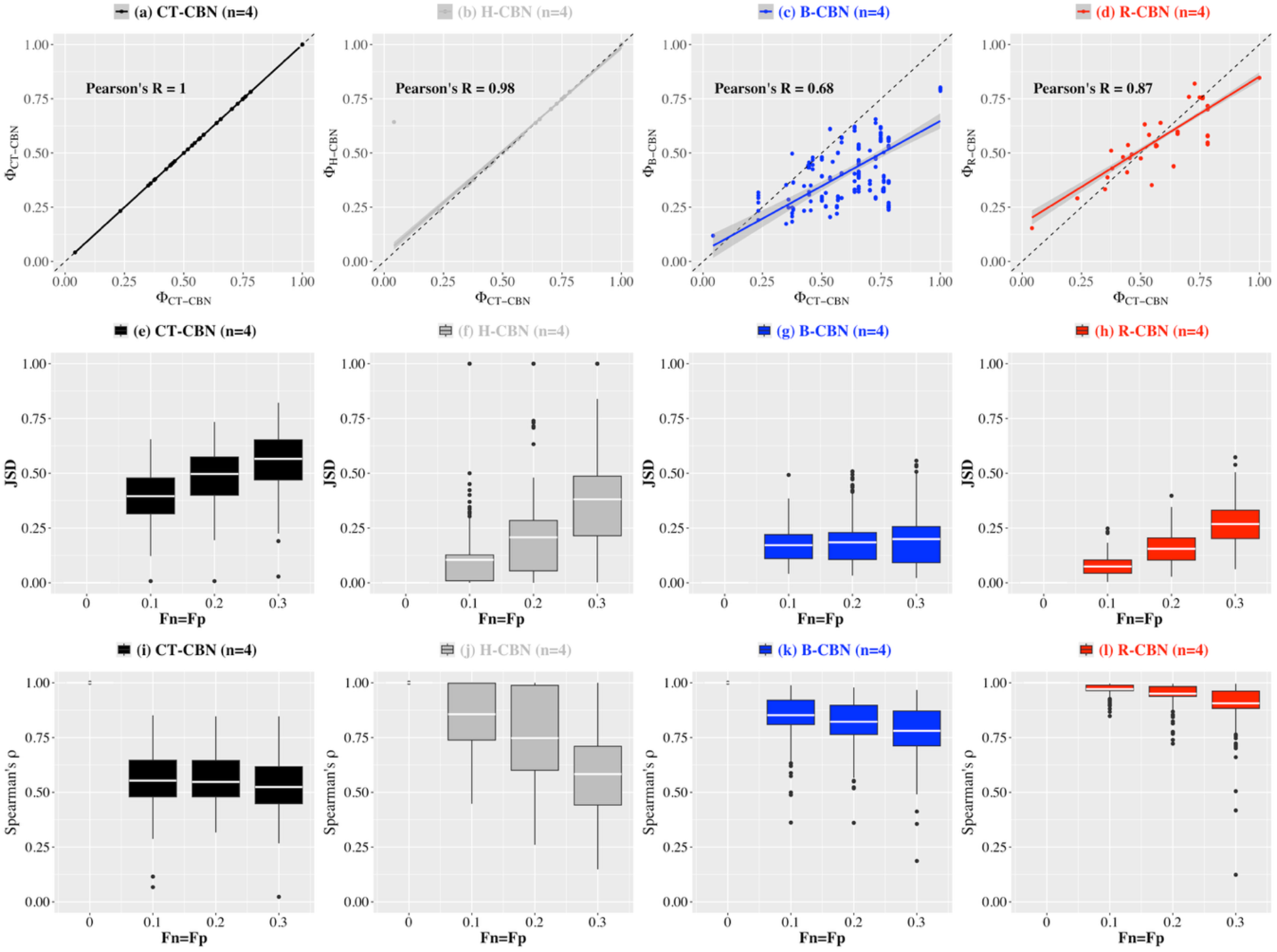
Benchmarking the CBN models (Synthetic Data including mutual exclusivity patterns; *n* = 4). In the upper panels, the horizontal axes indicate the predictability scores (Φ) quantified using CT-CBN model as the ground truth, and the vertical axes indicate the predictability scores (Φ) quantified in ideal conditions using 219 synthetic genotypic samples under a) CT-CBN, b) H-CBN, c) B-CBN and d) R-CBN. The box plots show the distribution of the Jensen-Shannon Divergence (*JSD*; the middle panels) and the Spearman’s correlation coefficient (*ρ*; the lower panels) between the pathway probability distributions before and after introducing genotypic errors of a given false positive (*Fp*) and false negative (*Fn*) rate (horizontal axes) using CT-CBN (panels e and i), H-CBN (panels f and j), B-CBN (panels g and k), and R-CBN (h and l). This figure is based on the second set of synthetic genotypic data in which patterns of mutual exclusivity are included in the genotype generation process. Note that the synthetic data analyses are based on pathways of length *n* = 4.

**Supplementary Figure S5.**
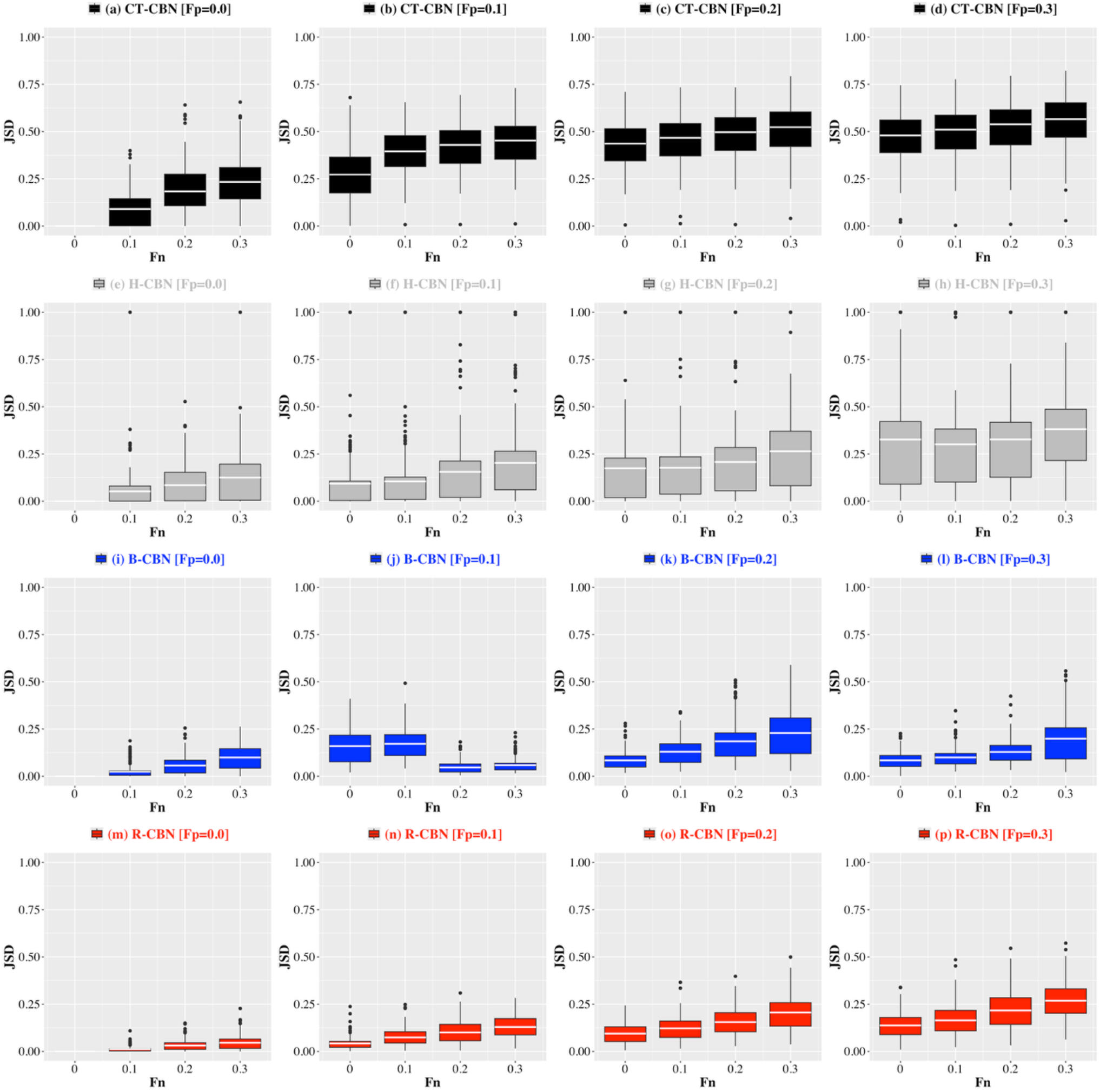
Robustness analysis I (Synthetic Data including mutual exclusivity patterns, *n* = 4 [All combinations of *Fn* and *Fp*]). The box plots show the distribution of the Jensen-Shannon Divergence (*JSD*) between the pathway probability distributions before and after introducing genotypic errors of a given false positive (*Fp*) rate (specified in the title of each panel) and false negative (*Fn*) rate (horizontal axes) using *i)* CT-CBN (black, the first row), *ii)* H-CBN (gray, the second row), *iii)* B-CBN (blue, the third row), and *iv)* R-CBN (red, the fourth row). This figure is based on the second set of synthetic genotypic data in which patterns of mutual exclusivity are included in the genotype generation process. Note that the synthetic data analyses are based on pathways of length *n* = 4.

**Supplementary Figure S6.**
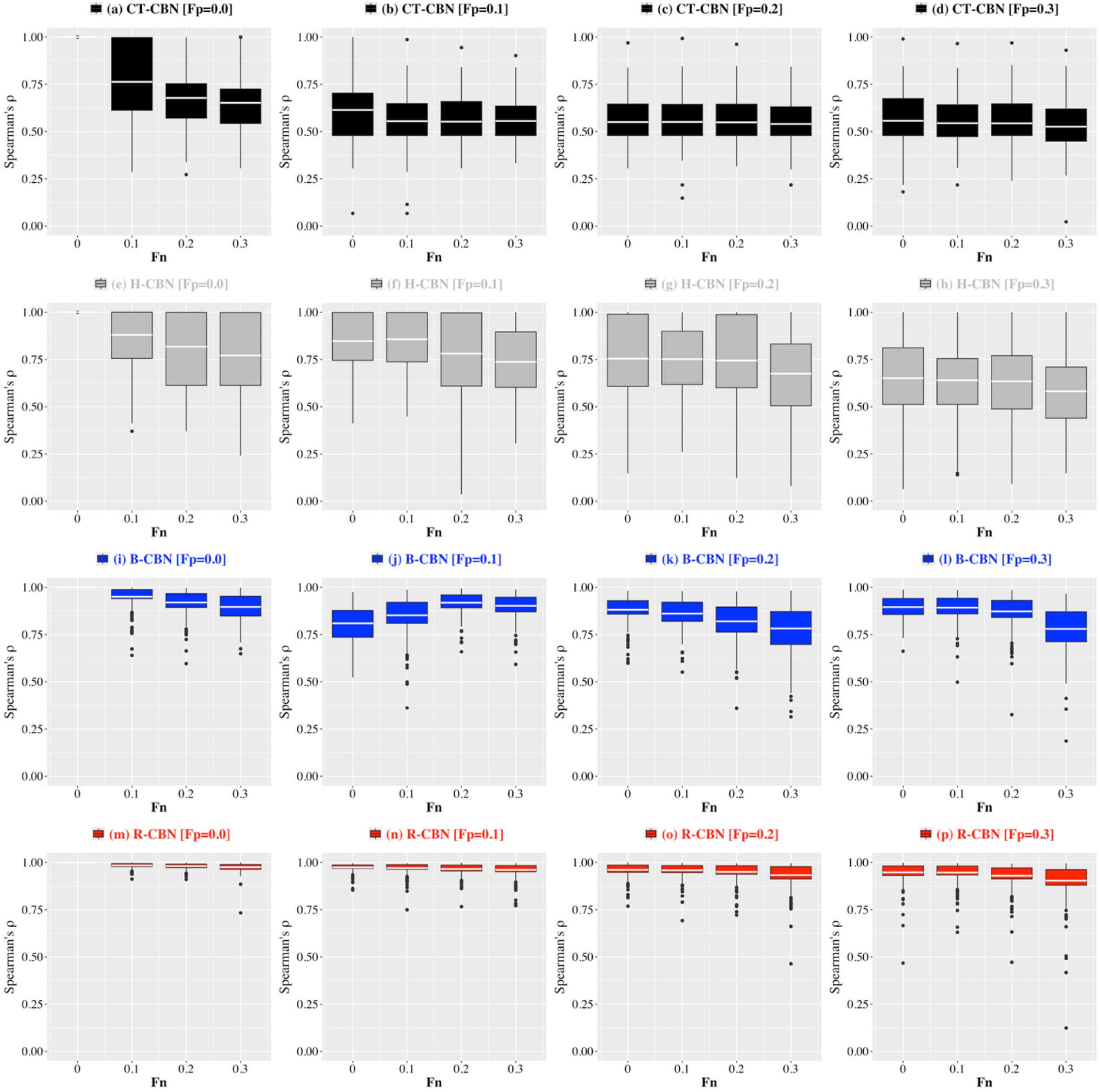
Robustness analysis II (Synthetic Data including mutual exclusivity patterns, *n* = 4 [All combinations of *Fn* and *Fp*]). The box plots show the distribution of the Spearman’s correlation coefficient (*ρ*) between the pathway probability distributions before and after introducing genotypic errors of a given false positive (*Fp*) rate (specified in the title of each panel) and false negative (*Fn*) rate (horizontal axes) using *i)* CT-CBN (black, the first row), *ii)* H-CBN (gray, the second row), *iii)* B-CBN (blue, the third row), and *iv)* R-CBN (red, the fourth row). This figure is based on the second set of synthetic genotypic data in which patterns of mutual exclusivity are included in the genotype generation process. Note that the synthetic data analyses are based on pathways of length *n* = 4.

**Supplementary Figure S7.**
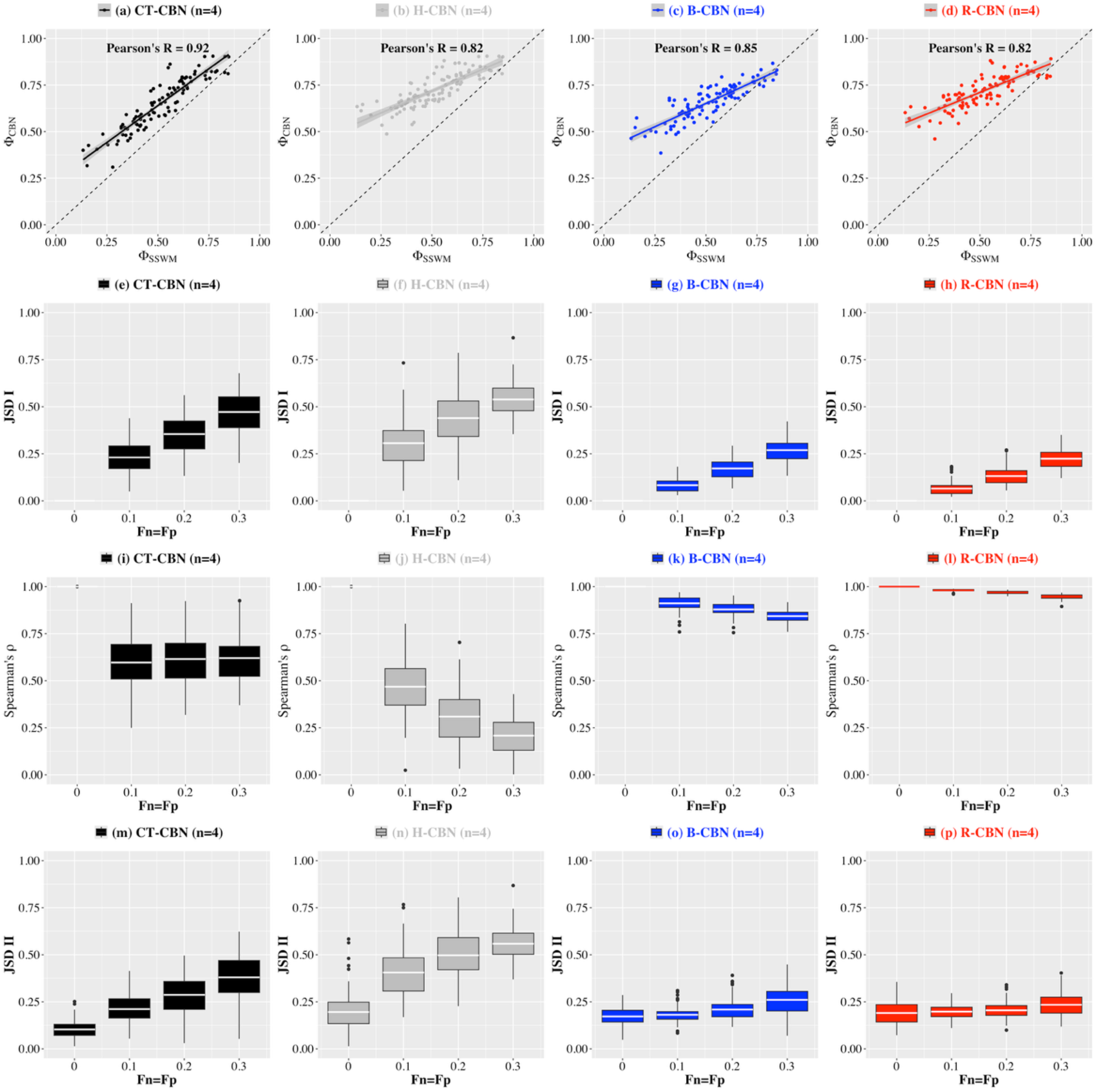
Benchmarking the CBN models (Simulated Data; *n* = 4 [Evolution under high mutation rates and fast detection regimes]). In the panels of the first row, the horizontal axes indicate the predictability scores (Φ_SSWM_) obtained using the pathway probability distributions inferred from 100 representable fitness landscape under the evolutionary SSWM-based model, while the vertical axes indicate the predictability scores (Φ_CBN_) obtained under a) CT-CBN, b) H-CBN, c) B-CBN and d) R-CBN models, which have been trained on the genotypic data resulting from evolutionary simulations with high mutation rates (10^−5^) and fast detection regimes (see Methods).The box plots show the distribution of the Jensen-Shannon Divergence (JDS I; the second row) and the Spearman’s correlation coefficient (*ρ*; the third row) between the pathway probability distributions before and after introducing genotypic errors of a given false positive (*Fp*) and false negative (*Fn*) rate (horizontal axes) using CT-CBN (panels *e* and *i*), H-CBN (panels *f* and *j*), B-CBN (panels *g* and *k*), and R-CBN (*h* and *l*). The box plots in the fourth row show the distribution of the Jensen-Shannon Divergence (*JDS II*) between the pathway probability distributions calculated using the SSWM evolutionary model and their corresponding CBN model (panels *m* to *p*). Note that this analysis is based on pathways of length *n* = 4, and so for each fitness landscape, 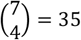 quartets are considered and the final metrics are averaged among them.

**Supplementary Figure S8.**
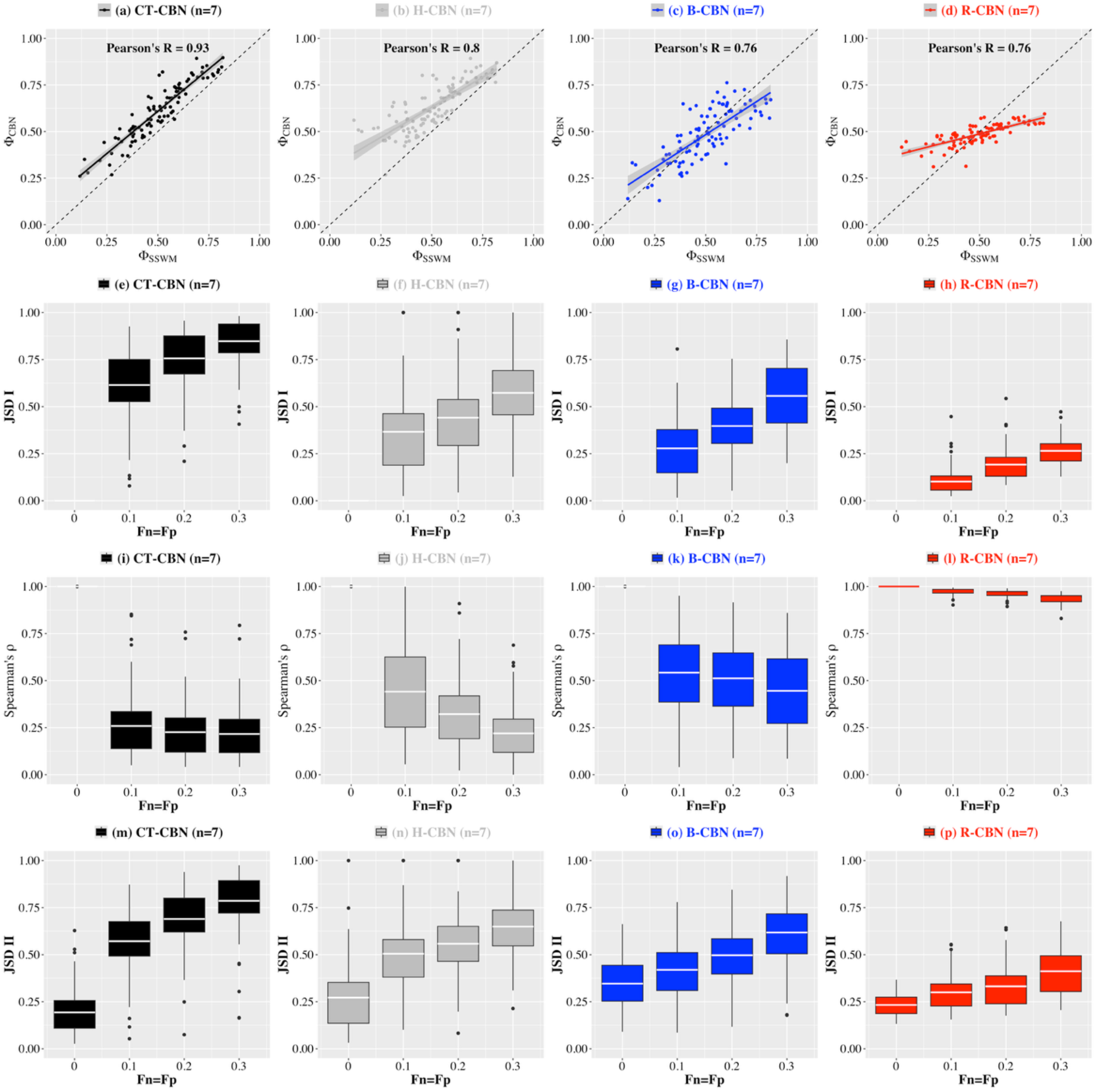
Benchmarking the CBN models (Simulated Data; *n* = 7 [Evolution under high mutation rates and fast detection regimes]). In the panels of the first row, the horizontal axes indicate the predictability scores (Φ_SSWM_) obtained using the pathway probability distributions inferred from 100 representable fitness landscape under the evolutionary SSWM-based model, while the vertical axes indicate the predictability scores (Φ_CBN_) obtained under a) CT-CBN, b) H-CBN, c) B-CBN and d) R-CBN models, which have been trained on the genotypic data resulting from evolutionary simulations with high mutation rates (10^−5^) and fast detection regimes (see Methods).The box plots show the distribution of the Jensen-Shannon Divergence (*JSD I*; the second row) and the Spearman’s correlation coefficient (*ρ*; the third row) between the pathway probability distributions before and after introducing genotypic errors of a given false positive (*Fp*) and false negative (*Fn*) rate (horizontal axes) using CT-CBN (panels *e* and *i*), H-CBN (panels *f* and *j*), B-CBN (panels *g* and *k*), and R-CBN (*h* and *l*). The box plots in the fourth row show the distribution of the Jensen-Shannon Divergence (*JSD II*) between the pathway probability distributions calculated using the SSWM evolutionary model and their corresponding CBN model (panels *m* to *p*). Note that this analysis is based on pathways of length *n* = 7.

## Notes

### Competing Interest Statement

The authors have declared no competing interest.

### Summary of Updates

I have substantially revised the original manuscript by adding a new dynamic programming algorithm to ensure the scalability of the inferential scheme. Consequently, I have added new analyses, new result sections, revised some of the previous figures and have added multiple new figures.

